# Telomere-to-telomere assembly of a complete human X chromosome

**DOI:** 10.1101/735928

**Authors:** Karen H. Miga, Sergey Koren, Arang Rhie, Mitchell R. Vollger, Ariel Gershman, Andrey Bzikadze, Shelise Brooks, Edmund Howe, David Porubsky, Glennis A. Logsdon, Valerie A. Schneider, Tamara Potapova, Jonathan Wood, William Chow, Joel Armstrong, Jeanne Fredrickson, Evgenia Pak, Kristof Tigyi, Milinn Kremitzki, Christopher Markovic, Valerie Maduro, Amalia Dutra, Gerard G. Bouffard, Alexander M. Chang, Nancy F. Hansen, Françoisen Thibaud-Nissen, Anthony D. Schmitt, Jon-Matthew Belton, Siddarth Selvaraj, Megan Y. Dennis, Daniela C. Soto, Ruta Sahasrabudhe, Gulhan Kaya, Josh Quick, Nicholas J. Loman, Nadine Holmes, Matthew Loose, Urvashi Surti, Rosa ana Risques, Tina A. Graves Lindsay, Robert Fulton, Ira Hall, Benedict Paten, Kerstin Howe, Winston Timp, Alice Young, James C. Mullikin, Pavel A. Pevzner, Jennifer L. Gerton, Beth A. Sullivan, Evan E. Eichler, Adam M. Phillippy

## Abstract

After nearly two decades of improvements, the current human reference genome (GRCh38) is the most accurate and complete vertebrate genome ever produced. However, no one chromosome has been finished end to end, and hundreds of unresolved gaps persist ^1,2^. The remaining gaps include ribosomal rDNA arrays, large near-identical segmental duplications, and satellite DNA arrays. These regions harbor largely unexplored variation of unknown consequence, and their absence from the current reference genome can lead to experimental artifacts and hide true variants when re-sequencing additional human genomes. Here we present a *de novo* human genome assembly that surpasses the continuity of GRCh38 ^2^, along with the first gapless, telomere-to-telomere assembly of a human chromosome. This was enabled by high-coverage, ultra-long-read nanopore sequencing of the complete hydatidiform mole CHM13 genome, combined with complementary technologies for quality improvement and validation. Focusing our efforts on the human X chromosome ^3^, we reconstructed the ∼2.8 megabase centromeric satellite DNA array and closed all 29 remaining gaps in the current reference, including new sequence from the human pseudoautosomal regions and cancer-testis ampliconic gene families (CT-X and GAGE). This complete chromosome X, combined with the ultra-long nanopore data, also allowed us to map methylation patterns across complex tandem repeats and satellite arrays for the first time. These results demonstrate that finishing the human genome is now within reach and will enable ongoing efforts to complete the remaining human chromosomes.

---

Complete, telomere-to-telomere reference assemblies are necessary to ensure that all genomic variants, large and small, are discovered and studied. Currently, unresolved regions of the human genome are defined by multi-megabase satellite arrays in the pericentromeric regions and the rDNA arrays on acrocentric short arms, as well as regions enriched in segmental duplications that are greater than hundreds of kilobases in length and greater than 98% identical between paralogs. Due to their absence from the reference, these repeat-rich sequences are often excluded from contemporary genetics and genomics studies, limiting the scope of association and functional analyses ^4,5^. Unresolved repeat sequences also result in unintended consequences such as paralogous sequence variants incorrectly called as allelic variants ^6^ and even the contamination of bacterial gene databases ^7^. Completion of the entire human genome is expected to contribute to our understanding of chromosome function ^8^ and human disease ^9^, and a comprehensive understanding of genomic variation will improve the driving technologies in biomedicine that currently use short-read mapping to a reference genome (e.g. RNA-seq ^10^, ChIP-seq ^11^, ATAC-seq ^12^).

The fundamental challenge of reconstructing a genome from many comparatively short sequencing reads—a process known as genome assembly—is distinguishing the repeated sequences from one another ^13^. Resolving such repeats relies on sequencing reads that are long enough to span the entire repeat or accurate enough to distinguish each repeat copy on the basis of unique variants ^14^. A recent *de novo* assembly of ultra-long (>100 kb) nanopore reads showed improved assembly continuity ^1^, but this proof-of-concept project sequenced the genome to only 5× depth of coverage and failed to assemble the largest human genomic repeats. Previous modeling based on the size and distribution of large repeats in the human genome predicted that an assembly of 30× ultra-long reads would approach the continuity of the human reference ^1^. Therefore, we hypothesized that high-coverage ultra-long-read nanopore sequencing would enable the first complete assembly of human chromosomes.

To circumvent the complexity of assembling both haplotypes of a diploid genome, we selected the effectively haploid CHM13hTERT cell line for sequencing (abbr. CHM13) ^15^. This cell line was derived from a complete hydatidiform mole with a 46,XX karyotype. The genomes of such molar pregnancies originate from a single sperm which has undergone post-meiotic chromosomal duplication and are, therefore, uniformly homozygous for one set of alleles. CHM13 has previously been used to patch gaps in the human reference ^2^, benchmark genome assemblers and diploid variant callers ^16^, and investigate human segmental duplications ^17^. Karyotyping of the CHM13 line confirmed a stable 46,XX karyotype, with no observable chromosomal anomalies (SFigs 1,2, SNote 1).

High molecular weight DNA from CHM13 cells was extracted and prepared for nanopore sequencing using a previously described ultra-long read protocol ^1^. In total, we sequenced 98 MinION flow cells for a total of 155 Gb (50× coverage, 1.6 Gb/flow cell, SNote 2). Half of all sequenced bases were contained in reads of 70 kb or longer (78 Gb, 25× genome coverage) and the longest validated read was 1.04 Mb. Once we had collected sufficient sequencing coverage for *de novo* assembly, we combined 39× of the ultra-long reads with 70× coverage of previously generated PacBio data ^18^ and assembled the CHM13 genome using Canu ^19^. This initial assembly totaled 2.90 Gbp with half of the genome contained in contiguous sequences (contigs) of length 75 Mbp or greater (NG50), which exceeds the continuity of the reference genome GRCh38 (75 vs. 56 Mbp NG50). Several chromosomes were captured in two contigs, broken only at the centromere (Fig 1a). The assembly was then iteratively polished by each technology in order of longest to shortest read lengths: Nanopore, PacBio, 10X Genomics / Illumina. Putative mis-assemblies were identified via analysis of independent linked-read sequencing (10X Genomics) and optical mapping (Bionano Genomics) data and the initial contigs broken at regions of low mapping coverage. The corrected contigs were then ordered and oriented relative to one another using the optical map and assigned to chromosomes using the human reference genome.

**Figure 1.**
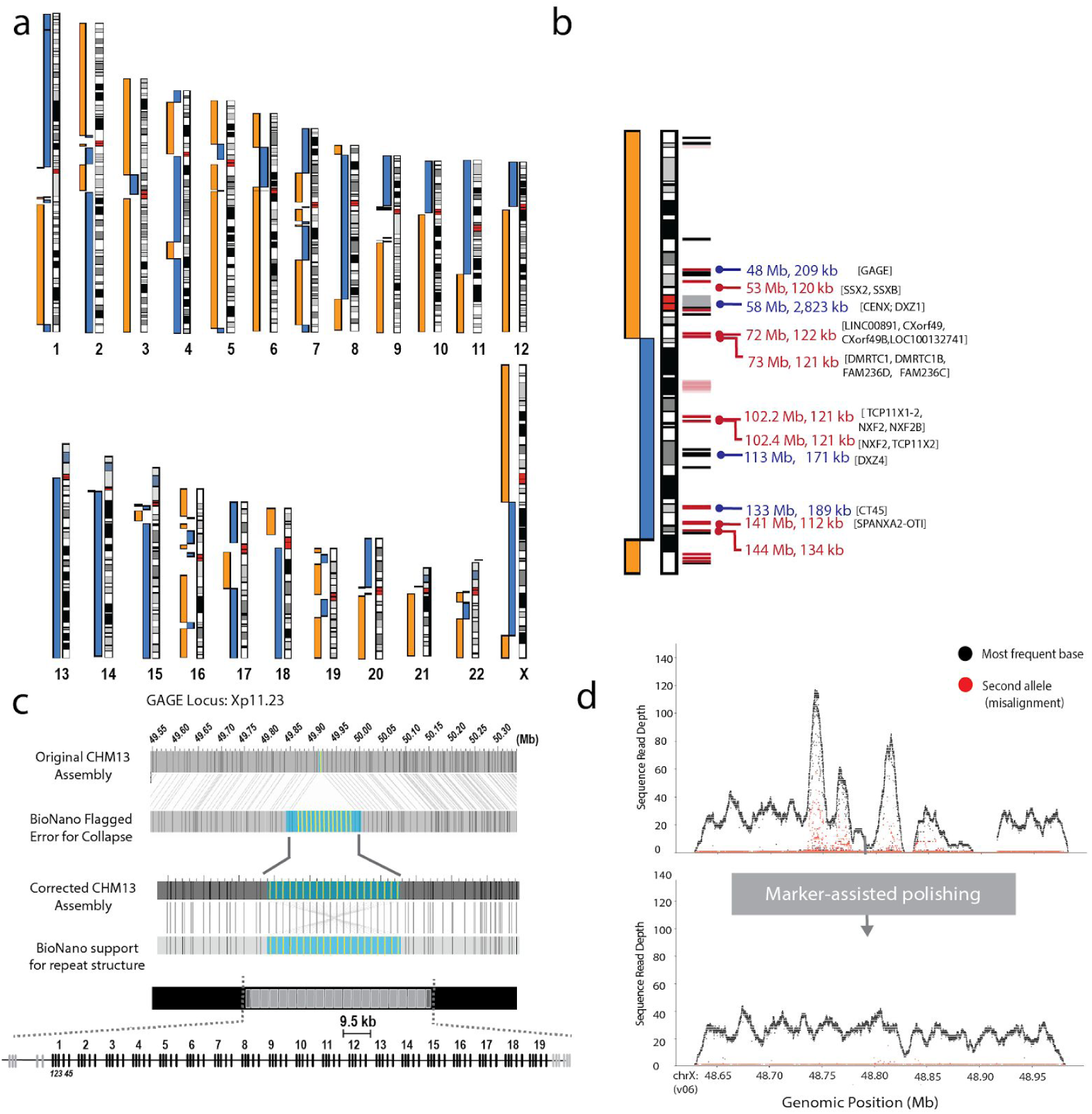
CHM13 whole-genome assembly and validation. (a) *De novo* assembly of the CHM13 genome using 39× of ultra-long Nanopore sequencing combined with 70× PacBio sequencing. Gapless contigs are illustrated as blue and orange bars next to the chromosome ideograms (two colors used only to better highlight contig breaks). Several chromosomes are broken only at centromeric regions (e.g. chr10, chr12, chr18, etc.). Large gaps between contigs (e.g. middle of chr1) indicate sites of large heterochromatic blocks or rDNA arrays where no GRCh38 reference sequence is available. (b) The X chromosome was selected for manual assembly, and was initially broken at three locations: the centromere (artificially collapsed in the assembly), a large segmental duplication (DMRTC1B, 120 kbp), and a second segmental duplication with a paralog on chromosome 2 (134 kbp). The relative placement of gaps in the GRCh38 reference are shown in the annotation track in black, known segmental duplications that are in red (with segmental duplications with paralogous sequence on the Y chromosome indicated in pink). Positions of repeats larger than 100 kb are indicated with the repeat sizing (kbs) in the CHM13 genome (blue, tandem repeats and red, segmental duplications). Tandem repeat classes, indicated in blue were resolved and evaluated by ddPCR and optical maps when applicable. (c) Mis-assembly of the GAGE locus identified by the optical map (top), and corrected version (bottom) showing the final assembly of 19 (9.5 kbp) full length repeat units and two partial repeats (d) Quality of the GAGE locus before and after polishing using unique (single-copy) markers to guide the correct placement of the long reads. Dots indicate coverage depth of the primary (black) and secondary (red) alleles recovered from mapped PacBio high-fidelity (HiFi) reads (SNote 3). Because the CHM13 genome is effectively haploid, regions of low coverage or increased secondary allele frequency indicate low-quality regions or potential repeat collapses. Marker-assisted polishing dramatically improved allele uniformity of across the entire GAGE locus.

**Table 1.** Assembly statistics for CHM13 and the human reference sorted by continuity

| Primary Technology | Assembly | Size (Gbp) | No. Ctgs | NG50 (Mbp) | %BACs resolved | BACs %idy all | BACs %idy uni |
| --- | --- | --- | --- | --- | --- | --- | --- |
| 56× 10x Genomics | Supernova (this paper) | 2.92 | 42,828 | 0.21 | 17.3 | 99.975 | 99.985 |
| 76× PacBio CLR | FALCON<sup>(57)</sup> | 2.88 | 1,916 | 28.2 | 36.37 | 99.981 | 99.995 |
| 24× PacBio HiFi | Canu<sup>(25)</sup> | 3.03 | 5,206 | 29.1 | 45.46 | 99.979 | 99.997 |
| Sanger BACs | GRCh38p13<sup>(2)</sup> | 3.11 | 1,590 | 56.4 | 85.63 | *99.731 | *99.768 |
| 39× Nanopore Ultra-Long | Canu (this paper) | 2.93 | 590 | 71.7 | 82.11 | 99.980 | 99.994 |

*Primary Technology:* sequencing technology used for contig assembly. The PacBio CLR assembly was additionally polished using Illumina data. The Nanopore Ultra-Long assembly was polished with the PacBio CLR and 10x Genomics data. GRCh38 is primarily based on Sanger-sequenced BACs, but has been continually curated and patched since the completion of the human genome project. *Assembly:* assembler used and reference to the published assembly. *Size:* sum of bases in the assembly in Gbp. GRCh38 assembly size includes 101 Mbp of alternative (ALT) sequences. *No. Ctgs:* total number of contigs in the assembly. *NG50:* half of the 3.09 Gbp human genome size contained in contigs of this length or greater in Mbp.
Supernova NG50 statistics were identical between the two reported pseudo-haplotypes. %BACs resolved: percentage of 341 “challenging” CHM13 BACs found intact in the assembly. BACs %idy all: median alignment accuracy versus all validation BACs. BACs %idy uni: median alignment accuracy versus the 31 validation BACs that occur outside of segmental duplications (SNote 3).

The final assembly consists of 2.94 Gbp in 590 contigs with a contig NG50 of 72 Mbp. We estimate the median consensus accuracy of this assembly to be >99.99%. Compared to other recent assemblies we resolved a greater fraction of the 341 CHM13 BAC sequences previously isolated and finished from segmentally duplicated and other difficult-to-assemble regions of the genome ^17^ (Tbl 1, SNote 3). Comparative annotation of our whole-genome assembly also shows a higher agreement of mapped transcripts than prior assemblies and only a slightly elevated rate of potential frameshifts compared to GRCh38 ^20^. Of the 19,618 genes annotated in the CHM13 *de novo* assembly, just 170 (0.86%) contain a predicted frameshift (STbl 1). When used as a reference sequence for calling structural variants in other genomes, CHM13 reports an even balance of insertion and deletion calls (SFig 3, SNote 4), as expected, whereas GRCh38 demonstrates a deletion bias, as previously reported ^21^. GRCh38 also reports more than twice the number of inversions than CHM13, suggesting that some mis-oriented sequences may remain in the current human reference. Thus, in terms of continuity, completeness, and correctness, our CHM13 assembly exceeds all prior human *de novo* assemblies, including the current human reference genome by some quality metrics (STbl 2).

Using this whole-genome assembly as a basis, we selected the X chromosome for manual finishing and validation due to its high continuity in the initial assembly, distinctive and well-characterized centromeric alpha satellite array ^3,8,22^, unique behavior during development ^23,24^, and disproportionate involvement in Mendelian disease ^3^. The *de novo* assembly of the X chromosome was broken in three places, at the centromere and two >100 kbp near-identical segmental duplications (Fig 1b). The two segmental duplications breaking the assembly were manually resolved by identifying ultra-long reads that completely spanned the repeats and were uniquely anchored on either side, allowing for a confident placement in the assembly. Assembly quality improvements of these difficult regions was evaluated by mapping an orthogonal set of PacBio high-fidelity (HiFi) long reads generated from CHM13 ^25^ and assessing read-depth over informative single nucleotide variant differences (Methods). In addition, experimental validation via droplet digital PCR (ddPCR) confirmed the now complete assembly correctly represents the tandem repeats of the CHM13 genome, including seven CT47 genes (7.02 ± 0.34), six CT45 genes (6.11 ± 0.38), 19 complete and two partial GAGE genes (19.9 ± 0.745), 55 DXZ4 repeats (55.4 ± 2.09), and a 2.8 Mbp centromeric DXZ1 array (1408 ± 40.69 2057 bp repeats) (Snote 5).

To assemble the X centromere, we constructed a catalog of structural and single-nucleotide variants within the ∼2 kbp canonical DXZ1 repeat unit ^26,27^ and used these variants as signposts ^8^ to uniquely tile ultra-long reads across the entire centromeric satellite array (DXZ1), as was previously done for the Y centromere ^28^. The DXZ1 array was estimated to be ∼2.8 Mbp (BglI, 2.87 +/- 0.16; BstEII, 2.82 +/- 0.03) by pulsed-field gel electrophoresis (PFGE) Southern blotting, wherein the resulting restriction profiles were in agreement with the structure of the predicted array assembly (Fig 2 a,b). Copy number estimates of the DXZ1 repeat by ddPCR were benchmarked against a panel of previously sized arrays by PFGE-Southern and provided further support for a ∼2.8 Mb array (1408 ± 81.38 copies of the canonical 2057 kbp repeat) (Fig 2c, STbl 3, SNote 6). Further, direct comparisons of DXZ1 structural variant frequency with PacBio HiFi data were highly concordant ^25^ (Fig 2d, SFig 4).

**Figure 2.**
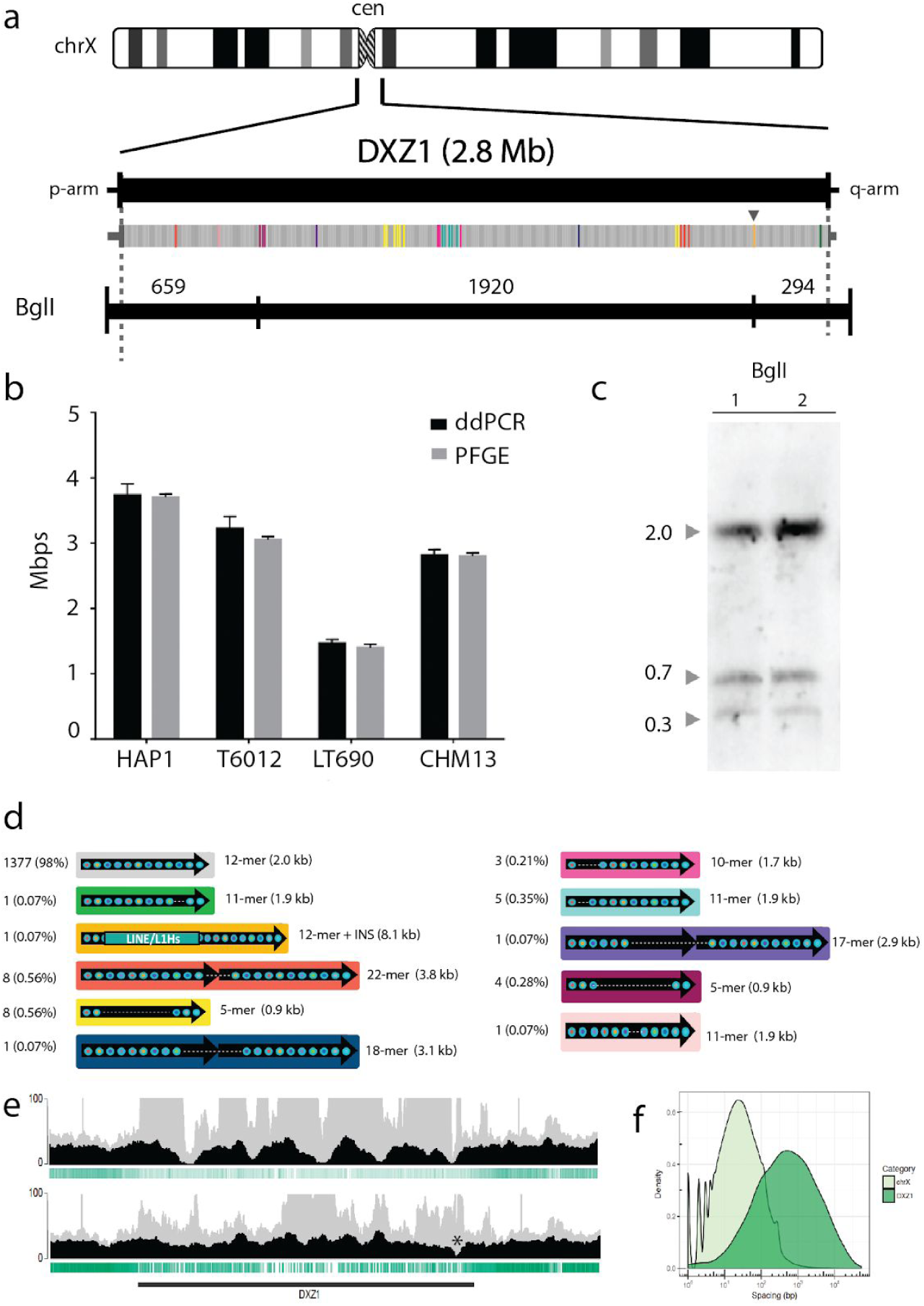
Validated structure of the 2.8 Mbp CHM13 X centromere array. (a) The reconstructed DXZ1 X centromere array is shown with all ∼2 kbp repeat units labeled by vertical bands, with grey indicating the canonical unit and colored bands indicating structural variants. A single LINE/L1Hs insertion was identified in the array, as marked by the arrowhead. Below, a predicted restriction map is shown for enzyme BglI, with dashed lines indicating regions outside of DXZ1 array that would be included in a restriction fragment. (b) Array size estimates were provided by ddPCR, which was optimized against PFGE Southerns of other available cell lines (HAP1, T6012, LT690). (c) Experimental PFGE Southern blotting is shown for a BglI digest in duplicate (band sizing indicated by triangles), that matches the *in silico* predicted band patterns (a) for the CHM13 array. (d) Catalog of 33 DXZ1 structural variants identified relative to the 2057 bp canonical repeat unit (grey), along with the number of instances observed, frequency in the array, number of alpha satellite monomers per repeat unit, and size of the variant repeat unit. (e) The black profile shows coverage depth of nanopore reads that could be uniquely anchored to the DXZ1 array before (top) and after (bottom) marker-assisted polishing (Methods). Single-copy, unique markers are shown as vertical green bands, with a decreased but non-zero density across the array. Coverage uniformity of the anchored reads improves after polishing, as well as the uniformity of reads mapped using standard Minimap2 mapping (light grey profile). One location of reduced coverage toward the right side of the array marks a possible issue remaining in the array (asterisk). (f) Distributions show the spacing between adjacent unique markers on chromosome X and DXZ1. On average, unique markers are found every 65 bases on chrX, but only every 22.8 kbp in DXZ1, with the longest gap between any two adjacent markers being 53 kbp.

Although unmatched in terms of continuity and structural accuracy, current long-read assemblies require rigorous consensus polishing to achieve maximum base call accuracy ^29–31^. Given the placement of each read in the assembly, these polishing tools statistically model the underlying signal data to make accurate predictions for each sequenced base. Key to this process is determining the correct placement of each read that will contribute to the polishing. Due to ambiguous read mappings, our initial polishing attempts actually decreased the assembly quality within the largest X chromosome repeats (SFig 5). To overcome this, we analyzed high-accuracy Illumina sequencing data to catalog short (21 bp), unique (single-copy) sequences present on the CHM13 X chromosome. Even within the largest repeat arrays, such as DXZ1, there was enough variation between repeat copies to induce unique 21-mer markers at semi-regular intervals (Fig 2 def, SFig 6). These markers were then used to inform the correct placement of long X-chromosome reads within the assembly (Methods). Using only high-confidence read mappings, two rounds of iterative polishing were performed for each technology, first with Oxford Nanopore ^32^, then PacBio ^29^, and finally 10X Genomics / Illumina ^33^, with consensus accuracy observed to increase after each round. Because the Illumina data was too short to confidently anchor, it was only used to polishing the unique regions of the chromosome where mappings were unambiguous. This detailed polishing process proved critical for accurately finishing X chromosome repeats that exceeded both Nanopore and PacBio read lengths.

Our manually finished X chromosome assembly is complete, gapless, and estimated to be at least 99.99% accurate (one error per 10 kbp, on average), which meets the original Bermuda Standards for finished genomic sequence ^34^. Accuracy is predicted to be slightly lower (median identity 99.3%) across the largest repeats, such as the DXZ1 satellite array, but this is difficult to measure due to a lack of BAC clones from these regions. Mapped long read and optical map data show uniform coverage across the completed X chromosome and no evidence of structural errors in regions that could be mapped (Fig 2e, SFig 7), while Strand-seq data confirm the absence of any inversion errors ^35,36^(SFig 8). Single nucleotide variant calling via long read mapping revealed lower initial assembly quality in the large, tandemly repeated GAGE and CT47 gene families, but these issues were resolved by polishing and validated via ultra-long read and optical mapping (Fig 1c,d, STbl 4, SFig 9, 10). A few isolated regions within the DXZ1 array show anomalous read coverage, which could be due to small structural errors beyond the resolution of PFGE-Southern or lower consensus quality in regions of the array containing relatively few unique markers needed for read anchoring and polishing (Fig 2 e-f, SFig 9, 10). Our complete telomere-to-telomere version of the X chromosome fully resolved 29 reference gaps ^3^, totaling 1,147,861 bp of previous N-bases.

Nanopore sequencing is sensitive to methylated bases as revealed by modulation to the raw electrical signal ^32,39^. Uniquely anchored ultra-long reads provide a new method to profile patterns of methylation over repetitive regions that are often difficult to detect with short-read sequencing. The X chromosome has many epigenomic features that are unique in the human genome. X-chromosome inactivation (XCI), in which one of the female X chromosomes is silenced early in development and remains inactive in somatic tissues, is expected to provide a unique methylation profile chromosome-wide. In agreement with previous studies ^40^, we observe decreased methylation across the majority of the pseudoautosomal regions (PAR1,2) located at both tips of the X chromosome arms (Fig 3a). The inactive X chromosome also adopts an unusual spatial conformation, and consistent with prior studies ^41,42^, CHM13 Hi-C data support two large superdomains partitioned at the macrosatellite repeat DXZ4 (SFig 11). On closer analysis of the DXZ4 array we found distinct bands of methylation (Fig 3c), with hypomethylation observed at the distal edge, which is generally concordant with previously described chromatin structure ^43^. Interestingly, we also identified a region of decreased methylation within the DXZ1 centromeric array (∼60 kbp, chrX:59,217,708–59,279,205, Fig 3b). To test if this finding was specific to the X array, or also found at other centromeric satellites, we manually assembled a ∼2.02 Mbp centromeric array on chromosome 8 (D8Z2) ^44,45^ and employed the same unique marker mapping strategy to confidently anchor long reads across the array. In doing so, we identified another hypomethylated region within the D8Z2 array, similar to our observation on the DXZ1 array (SFig 12), further demonstrating the capability of our ultra-long read mapping strategy to provide base-level chromosome-wide DNA methylation maps. Future studies will be needed to evaluate the potential importance, if any, of these methylation patterns.

**Figure 3.**
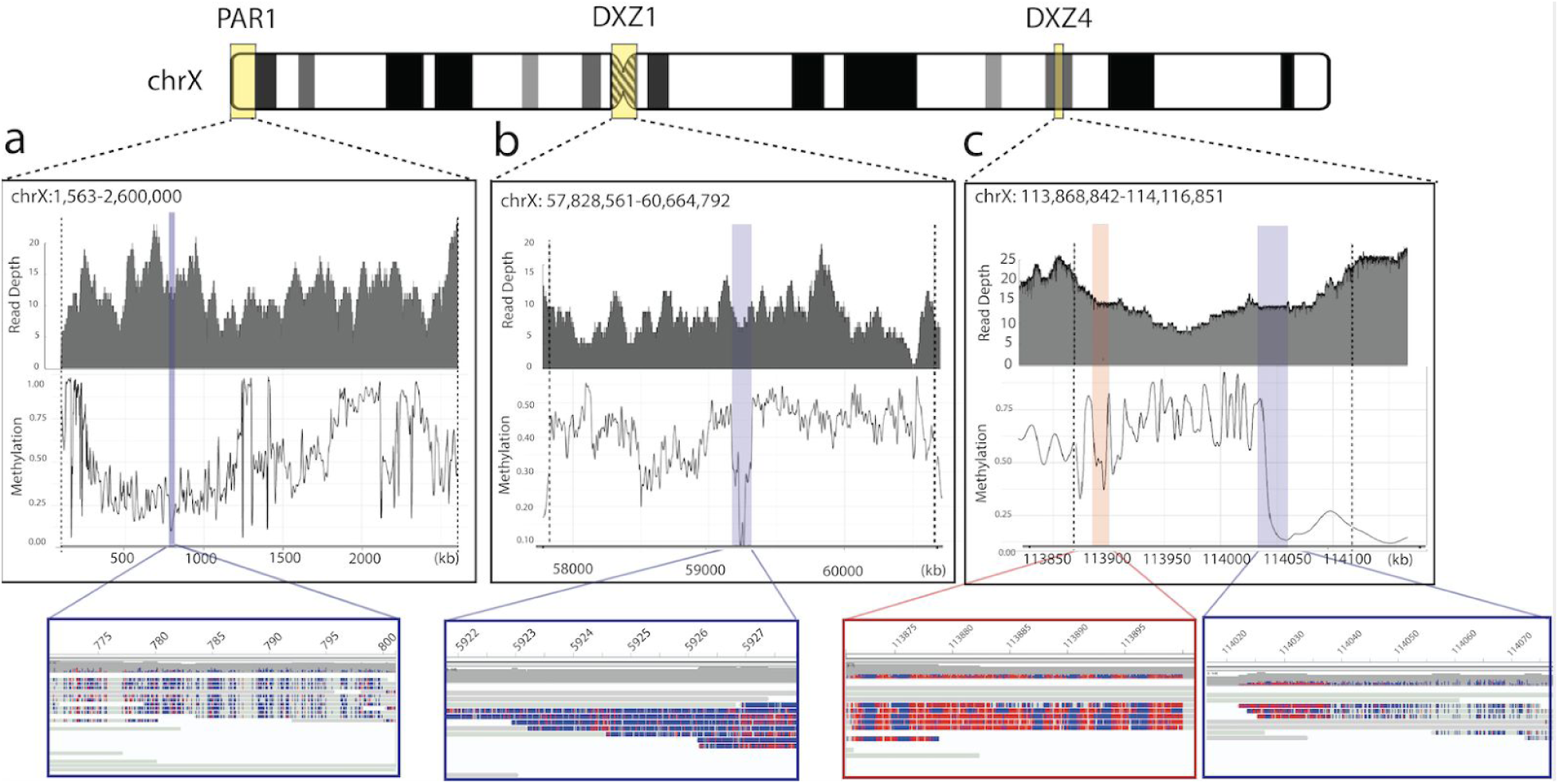
Chromosome-wide analysis of CpG methylation. Methylation estimates were calculated by smoothing methylation frequency data with a window size of 500 nucleotides. Coverage depth and high quality methylation calls (|log-likelihood| > 2.5) for PAR1, DXZ1, and DXZ4 are shown as insets. Only reads with a confident unique anchor mapping and the presence of at least one high-quality methylation call were considered. (a) Nanopore coverage and methylation calls for the pseudoautosomal region 1 (PAR1) of chromosome X (1,563–2,600,000). Bottom IGV inset shows a region of hypomethylation within PAR1 (770,545–801,293) with unmethylated bases in blue and methylated bases in red. (b) Methylation in the DXZ1 array, with bottom IGV inset showing a ∼60 kbp region of hypomethylation near the centromere of chromosome X (59,217,708–59,279,205). (c) Vertical black dashed lines indicate the beginning and end coordinates of the DXZ4 array. Left IGV inset shows a methylated region of DXZ4 in chromosome X (113,870,751–113,901,499), and right IGV inset shows a transition from a methylated to unmethylated region of DXZ4 (114,015,971–114,077,699).

This first complete telomere-to-telomere assembly of a human chromosome demonstrates that it may now be possible to finish the entire human genome using available technologies. Important challenges remain going forward. Applying these approaches, for example, to diploid samples will require phasing the underlying haplotypes to avoid mixing regions of complex structural variation. Our preliminary analysis of other chromosomes shows that regions of duplication and centromeric satellites larger than that of the X chromosome will require additional methods development. This is especially true of the acrocentric human chromosomes whose massive satellite and segmental duplications have yet to be resolved at the sequence level. Although we have focused here on finishing the X chromosome, our whole-genome assembly has reconstructed several other chromosomes with only a few remaining gaps and can serve as the basis for completing additional human chromosomes (Fig 1). Efforts to finally complete the human reference genome will help advance the necessary technology towards our ultimate goal of telomere-to-telomere assemblies for all human genomes.

## Methods

### Cell culture

Cells from a case of a complete hydatidiform mole CHM13 were cultured, karyotyped using Q banding and cryopreserved at Magee-Womens Hospital (Pittsburgh, PA). The thawed cells were subsequently immortalized using Human telomerase reverse transcriptase (hTERT). The CHM13 cells were cultured in complete AmnioMax C-100 Basal Medium (ThermoFisher Scientific, Carlsbad, CA) supplemented with 1% Penicillin-Streptomycin (ThermoFisher Scientific, Carlsbad, CA) and grown in a humidity-controlled environment at 37°C, with 95% O_2_ and 5% CO_2_. Fresh medium was exchanged every 3 days and all cells used for this study did not exceed passage 10.

### Karyotyping

Metaphase slide preparations were made from human hydatidiform mole cell line CHM13, and prepared by standard air-drying technique as previously described ^46^. DAPI banding techniques were performed to identify structural and numerical chromosome aberrations in the karyotypes according to the ISCN ^47^. Karyotypes were analyzed using a Zeiss M2 fluorescence microscope and Applied Spectral Imaging software (Carlsbad, CA).

### DNA extraction, library preparation, and sequencing

High-molecular-weight (HMW) DNA was extracted from 5×107 CHM13 cells using a modified Sambrook and Russell protocol ^1,48^. Libraries were constructed using the Rapid Sequencing Kit (SQK-RAD004) from Oxford Nanopore Technologies with 15 μg of DNA. The initial reaction was typically divided into thirds for loading and FRA Buffer (104 mM Tris pH 8.0, 233 mM NaCl) was added to bring the volume to 21 ul. These reactions were incubated at 4°C for 48 hrs to allow the buffers to equilibrate before loading. Most sequencing was performed on the Nanopore GridION with FLO-MIN106 or FLO-MIN106D R9 flow cells, with the exception of one Flongle flow cell used for testing. Sequencing reads used in the initial assembly were first basecalled on the sequencing instrument. After all data was collected, the reads were basecalled again using the more recent Guppy algorithm (version 2.3.1 with the “flip-flop” model enabled).

A 10X Genomics linked-read genomic library was prepared from 1 ng of high molecular weight genomic DNA using a 10X Genomics Chromium device and Chromium Reagent Kit v2 according to the manufacturer’s protocol. The library was sequenced on an Illumina NovaSeq 6000 DNA sequencer on an S4 flow cell, generating 586 million paired-end 151 base reads. The raw data was processed using RTA3.3.3 and bwa0.7.12 ^49^. The resulting molecule size was calculated to be 130.6 kb from a Supernova ^50^ assembly.

DNA was prepared using the ‘Bionano Prep Cell Culture DNA Isolation Protocol’. After cells were harvested, they were put through a number of washes before embedding in agarose. A proteinase K digestion was performed, followed by additional washes and agarose digestion. The DNA was assessed for quantity and quality using a Qubit dsDNA BR Assay kit and CHEF gel. A 750 ng aliquot of DNA was labeled and stained following the Bionano Prep Direct Label and Stain (DLS) protocol. Once stained, the DNA was quantified using a Qubit dsDNA HS Assay kit and run on the Saphyr chip.

Hi-C libraries were generated, in replicate, by Arima Genomics using four restriction enzymes. After the modified chromatin digestion, digested ends were labelled, proximally ligated, and then proximally-ligated DNA was purified. After the Arima-HiC protocol, Illumina-compatible sequencing libraries were prepared by first shearing then size-selecting DNA fragments using SPRI beads. The size-selected fragments containing ligation junctions were enriched using Enrichment Beads provided in the Arima-HiC kit, and converted into Illumina-compatible sequencing libraries using the Swift Accel-NGS 2S Plus kit (P/N: 21024) reagents. After adapter ligation, DNA was PCR amplified and purified using SPRI beads. The purified DNA underwent standard QC (qPCR and Bioanalyzer) and sequenced on the HiSeq X following manufacturer’s protocols.

### Whole-genome assembly

#### Nanopore and PacBio whole-genome assembly

Canu v1.7.1 ^19^ was run with all rel1 Oxford Nanopore data (on-instrument basecaller, rel1) generated on or before 2018/11/07 (totaling 39x coverage) and PacBio sequences (SRA: PRJNA269593) generated in 2014 and 2015 (totaling 70x coverage) ^2^,18.

### Manual gap closure

Gaps on the X chromosome were closed by mapping all reads against the assembly and manually identifying reads joining contigs that were not included in the automated Canu assembly. This generated an initial candidate chromosome assembly, with the exception of the centromere. Four regions of the candidate assembly were found to be structurally inconsistent with the Bionano optical map and were corrected by manually selecting reads from those regions and locally reassembling with Canu ^19^ and Flye v2.4 ^51^. Low coverage long reads that confidently spanned the entire repeat region were used to guide and evaluate the final assembly where available. Evaluation of copy number and repeat organization between the re-assembled version and spanning reads was performed using HMMER (v3) ^52,53^ trained on a specific tandem repeat unit, and the reported structures were manually compared. Default parameters for Minimap2 ^54^ resulted in uneven coverage and polishing accuracy over tandemly repeated sequences. This was successfully addressed by increasing the Minimap2 -r parameter from 500 to 10000 and increasing the maximum number of reported secondary alignments (-N) from 5 to 50. Final evaluation of repeat base-level quality was determined by mapping of PacBio datasets (CLR and HiFi) (SFigs 5,9,10, SNote 3).

The alpha satellite array in the X centromere, due to its availability as a haploid array in male genomes, is one of the best studied centromeric regions at the genomic level, with a well-defined 2 kbp repeat unit ^26^, physical/genetic maps ^8,55^ and an expected range of array lengths ^22^. We initially generated a database of alpha satellite containing ultra-long reads, by labeling those reads with at least one complete consensus sequence ^27^ of a 171 bp canonical repeat in both orientations, as previously described ^56^. Reads containing alpha in the reverse orientation were reverse complemented, and screened with HMMER (v3) using a 2057 bp DXZ1 repeat unit. We then employed run-length encoding in which runs of the 2057 bp canonical repeat (defined as any repeat in the range of min: 1957 bp, max: 2157 bp) were stored as a single data value and count, rather than the original run. This allowed us to redefine all reads as a series of variants, or repeats that differ in size/structure from the expected canonical repeat unit, with a defined spacing in between. Identified CHM13 DXZ1-SVs in the UL-read data were compared to a library of previously characterized rearrangements in published PacBio (CLR ^18,57^ and HiFi ^25^) using alpha-CENTUARI, as described ^56^. Output annotation of SVs and canonical DXZ1 spacing for each read were manually clustered to generate six initial contigs, two of which are known to anchor into the adjacent Xp or Xq. To define the order and overlap between contigs, we identified all 21-mers that had an exact match within the high-quality DXZ1 array data obtained from CRISPR-Cas9 duplexSeq targeted resequencing ^58^ (SNote 7). Overlap between the two or more 21-mers with equal spacing guided the organization of the assembly. Orthogonal validation of the spacing between contigs (and contig structure) was supported with additional ultra-long read coverage, providing high-confidence in repeat unit counts for all but three regions.

### Chromosome X long-read polishing

We used a novel mapping pipeline to place reads within repeats using unique markers. *k*-mers were collected from the 10x Genomics / Illumina dataset, after trimming off the barcodes (the first 23 bases of the first read in a pair). The read was placed in the location of the assembly having the most unique markers in common with the read. Alignments were further filtered to exclude short and low identity alignments. This process was repeated after each polishing round, with new unique markers and alignments recomputed after each round. Polishing proceeded with one round of Racon followed by two rounds of Nanopolish and two rounds of Arrow. Post-polishing, all previously flagged low-quality loci showed significant improvement, with the exception of 138.6–139.7 which still had a coverage drop and was replaced with an alternate patch assembly generated by Canu using PacBio HiFi data.

### Whole-genome long-read polishing

The rest of the whole-genome assembly was polished similarly to the X chromosome, but without the use of unique *k*-mer anchoring. Instead, two rounds of Nanopolish, followed by two rounds of Arrow, were run using the above parameters, which rely on the mapping quality and length / identity thresholds to determine the best placements of the long reads. As no concerted effort was made to correctly assemble the large satellite arrays on chromosomes outside of the X, this default polishing method was deemed sufficient for the remainder of the genome. However, future efforts to complete these remaining chromosomes are expected to benefit from the unique *k*-mer anchoring mapping approach.

### Whole-genome short-read polishing

The 10x Genomics / Illumina data was used for a final polishing of the whole assembly, including the X chromosome, but using only unambiguous mappings and allowing only indel corrections (SNote 3).

### Methylation analysis

To measure CpG methylation in nanopore data we used Nanopolish ^32^. Nanopolish employs a Hidden Markov Model (HMM) on the nanopore current signal to distinguish 5-methylcytosine from unmethylated cytosine. The methylation caller generates a log-likelihood value for the ratio of probability of methylated to unmethylated CGs at a specific *k*-mer. We next filtered methylation calls using the nanopore_methylation_utilities tool (https://github.com/isaclee/nanopore-methylation-utilities), which uses a log-likelihood ratio of 2.5 as a threshold for calling methylation ^59^. CpG sites with log-likelihood ratios greater than 2.5 (methylated) or less than −2.5 (unmethylated) are considered high-quality and included in the analysis. Reads that do not have any high-quality CpG sites are filtered from the bam for subsequent methylation analysis. Figure 3 shows coverage of reads with at least one high quality CpG site. Nanopore_methylation_utilities integrates methylation information into the alignment BAM file for viewing in IGV’s ^60^ bisulfite mode and also creates Bismark-style files which we then analyzed with the R Bioconductor package BSseq ^61^. We used the BSmooth algorithm ^61^ within the BSseq package for smoothing the data to estimate the methylation level at specific regions of interest.

### Data availability

Original data generated at SIMR that underlies this manuscript can be accessed from the Stowers Original Data Repository at http://www.stowers.org/research/publications/libpb-1453. Genome assemblies and sequencing data including raw signal files (FAST5), event-level data (FAST5), base-calls (FASTQ), and alignments (BAM/CRAM) are available as an Amazon Web Services Open Data set. Instructions for accessing the data, as well as future updates to the raw data and assembly, are available from https://github.com/nanopore-wgs-consortium/chm13. All data is additionally archived and available under NCBI BioProject accession PRJNA559484 (*submission in progress*) and NCBI accession ### for the whole-genome assembly and completed X chromosome.

## Acknowledgements

We acknowledge helpful conversations with Isac Lee on methylation analysis, and a helpful review of the manuscript by Huntington F. Willard. Funding support: NIH/NHGRI R21 1R21HG010548-01 (KHM); Intramural Research Program of the National Human Genome Research Institute, National Institutes of Health (SK, AR, VM, AD, GGB, AMC, NFH, AY, JCM, AMP); Korea Health Technology R&D Project through the Korea Health Industry Development Institute HI17C2098 (AR); Intramural Research Program of the National Library of Medicine, National Institutes of Health (VAS, FT-N); Common Fund, Office of the Director, NIH (VM); Stowers Institute for Medical Research (EH, TP, JLG); NIH R01 GM124041 (BAS); NIH HG002385 and HG010169 (EEE); EEE is an investigator of the Howard Hughes Medical Institute; National Library of Medicine Big Data Training Grant for Genomics and Neuroscience 5T32LM012419-04 (MRV); NIH 1F32GM134558-01 (GAL); NIH/NHGRI U54 1U54HG007990 and W. M. Keck Foundation DT06172015 and NIH/NHLBI U01 1U01HL137183 and NIH/NHGRI/EMBL 2U41HG007234 (BP); NIH/NHGRI R01 HG009190 and NIGMS T32 GM007445 (WT, AG); NIH R01CA181308 (RAR); NIH/NHGRI 2R44HG008118 (ADS and SS); Wellcome Trust [212965/Z/18/Z] (NH, NJL, ML); National Institute for Health Research (NIHR) Surgical Reconstruction and Microbiology Research Centre (SRMRC) (JQ). The views expressed are those of the author(s) and not necessarily those of the NIHR or the Department of Health and Social Care. This work utilized the computational resources of the NIH HPC Biowulf cluster (https://hpc.nih.gov).

## Author Contributions

SB, GAL, KT, VM, GGB, MYD, DCS, RS, GK, NH, ML, AY, JCM, EEE performed CHM13 nanopore sequencing, cell line preparation, and primary data analysis. AY and JCM generated 10x whole genome sequencing and assembly. BAS performed PFGE-Southern blotting array size analysis. MK, CM, RF, TAGL, and IH generated bionano data and performed data analysis. JF and RR performed CRISPr-DS analysis. EH, TP, JLG performed ddPCR and SKY analysis. EP, AD, EH, TP, JLG performed CMH13 cell line karyotyping. KHM performed repeat characterization and satellite DNA assembly. KHM, SK, MRV, AMC, and AMP performed automated and manual assembly. KHM, SK, AR, MRV, GAL, DP, JW, WC, KH, EEE, and AMP performed assembly curation and validation. SK, AR, and AMP performed marker-based assembly polishing. AG and WT performed methylation analysis. AB and PAP generated automated satellite DNA assemblies. ADS and JMB and SS performed Hi-C CHM13 sequencing. AR performed Hi-C analysis. NFH performed SV analysis. JA and BP performed annotation analysis. VAS and FTN performed alignment versus RefSeq, repeat characterization, and frameshift analysis. US provided access to critical resources. JQ developed the initial ultra-long read protocol and updated to current chemistry. NJL provided an Amazon Web Services (AWS) account and coordinated data sharing. KHM, SK, AR, MRV, and AMP developed figures. KHM and AMP coordinated the project. KHM, SK, and AMP drafted the manuscript. All authors read and approved the final manuscript.

