## Supplemental Material for "Telomere-to-telomere assembly of a complete human X chromosome"

**This PDF file includes:**
**Supplemental Notes 1 to 7**
**Figures S1 to S12**
**Tables S1 to S4**
**References**

#### Supplementary Note 1. CHM13 cell line and chromosome characterization

##### CHM13hTERT Cell Line

CHM13 cells were originally grown in culture from one case of a hydatidiform mole at Magee-Womens Hospital (Pittsburgh, PA) as part of a research study (IRB MWH-20-054). Cryogenically frozen cells from this culture were grown and transformed using human telomerase reverse transcriptase (TERT) to develop a cell line. This cell line retains a 46,XX karyotype and complete homozygosity.

##### Spectral karyotyping (SKY)

Spectral imaging was performed using In laser-scanning confocal microscopes LSM-710 and LSM-780 (Carl Zeiss Microimaging, Jena, Germany). Both microscopes were equipped with a QUASAR detection unit that can acquire with a single scan an entire range of emission wavelengths (in 10 nm increments) for subsequent spectral unmixing. For spectral imaging, 3 excitation laser lines were utilized: 488, 561, and 633 nm. Images were collected with 3 different dichroics: the first passing 488 nm excitation, the second passing 488 nm and 561 nm excitation, and the third passing all 3 laser lines. In addition, a 405 nm laser was used to acquire a Hoechst 33342– stained DNA image for segmentation, with emission collected at ~450 nm. All images were acquired with either a 40× or 63× Plan Apochromat objective (Carl Zeiss Microimaging). Pinhole settings were optimized for background reduction and signal-to-noise ratio. Image processing and karyotyping of the CHM13 line were performed with a set of custom open source ImageJ (NIH, Bethesda, MD) plugins called Karyotype Identification via Spectral Separation (KISS), freely available at http://research.stowers.org/imagejplugins/KISS_analysis.html. Briefly, the plugins perform interactive background subtraction, spectral unmixing, interactive chromosome segmentation, and interactive karyotyping based on dye composition. Chromosome segmentation is performed using a semi-automated method based on the Hoechst image. First, the image is smoothed with a Gaussian blur with a 1 pixel standard deviation and then segmented with a manually chosen fractional threshold and object area limits to eliminate dirt and intact nuclei. Next, chromosomes too close to be separated by thresholding are manually separated. Finally overlapping chromosomes are separated into non-overlapping parts and then linked together for karyotyping. A total of 10 SKY images were evaluated to assess the stability of the CHM13 line.

#### Supplementary Note 2. Library preparation and sequencing

**Oxford Nanopore**

Library preparation and nanopore sequencing was performed as previously described [1], with the following updates. Generation of ultra-long reads employs the Rapid Sequencing Kit (Oxford Nanopore Technologies, UK) and comprises two steps: tagmentation of DNA by a transposase complex followed by attachment of the sequencing adapter. Previous work was performed using kit SQK-RAD002 which was replaced by SQK-RAD003 in Jun 2017. Testing performed on this kit indicated difficulty generating ultra-long reads was due to a protocol change which doubled the standard input required from 200 ng to 400 ng and a reformulation of the FRM reagent (now called FRA). This protocol resulted in low efficiency libraries when using HMW DNA input. Testing showed that reducing the volume of fragmentation reagent from 5 ul to 1.5 ul and the addition of 0.02% Triton-X100 final concentration could restore library performance. The modifications are included in the 'Ultra-long read sequencing protocol for RAD004' (dx.doi.org/10.17504/protocols.io.mrxc57n) used here.

High-molecular-weight genomic DNA from the CHM13hTERT cell line was obtained using a modified Sambrook and Russell DNA extraction method before preparing ultra-long read sequencing libraries using the protocol above. Briefly, 16 µl of DNA from the Sambrook extraction at approximately 1 µg/µl, manipulated with a wide-bore P20 pipette tip, was placed in a 0.2 ml PCR tube, with 1 µl removed to confirm quantification value. 3.5ul EB and 1.5 µl FRA (SQK-RAD004, ONT) was added and mixed slowly ten times by gentle pipetting with a wide-bore pipette tip moving only 18 µl. After mixing, the sample was incubated at 30 °C for 1 min followed by 80 °C for 1 min on a thermocycler. After this, 1 µl RAP (SQK-RAD004, ONT) was added and mixed slowly ten times by gentle pipetting with a cut-off pipette tip moving only 14 µl. The library was then incubated at room temperature for 30 min to allow adapter attachment. Libraries are divided, diluted and incubated for 48 hours (as discussed in updates above). To load the library, 34 µl SQB (SQK-RAD004, ONT) was mixed with 20 µl nuclease-free water, and this was added to the library. Using a P100 wide-bore tip set to 75 µl, this library was mixed by pipetting slowly five times. This extremely viscous sample was loaded onto the "spot on" port and entered the flow cell by capillary action. The standard loading beads were omitted from this protocol owing to excessive clumping when mixed with the viscous library.

GridION sequencing was performed as per manufacturer's guidelines using R9/R9.4 flow cells (FLO-MIN106 or FLO-MIN106D, ONT), and controlled using Oxford Nanopore Technologies MinKNOW (version 3.4.5) software. The specific versions of the software used varied from run to run but can be determined by inspection of the provided fast5 files. This generated the rel1 dataset.

Reads from all sites were copied to the NIH Biowulf HPC cluster, where base calling was performed using Guppy (flip-flop version 2.3.1) to generate the updated dataset (referred to as rel2).

**10x Genomics**
A linked read genomic library was prepared from one nanogram of high molecular weight genomic DNA using a 10x Genomics Chromium device and Chromium Reagent Kit v2 according to manufacturer's protocol. The library was sequenced on a NovaSeq 6000 DNA sequencer (Illumina, Inc.) on an S4 flow cell, generating 586M paired-end 151 base reads. The raw data was processed using RTA3.3.3 and bwa0.7.12. The resulting molecule size was calculated to be 130.6 kb from a Supernova assembly.

**Bionano optical mapping**
DNA was prepared using the 'Bionano Prep Cell Culture DNA Isolation Protocol'. After cells were harvested, they were put through a number of washes before embedding in agarose. A proteinase K digestion was performed, followed by additional washes and agarose digestion. From this point, the DNA was drop dialyzed and allowed to equilibrate at room temperature for two days. The DNA was assessed for quantity and quality using a Qubit dsDNA BR Assay kit and CHEF gel. A 750 ng aliquot of DNA was labeled and stained following the Bionano Prep Direct Label and Stain (DLS) protocol. Once stained, the DNA was quantified using a Qubit dsDNA HS Assay kit and run on the Saphyr chip.

**Hi-C sequencing**

Hi-C libraries were generated, in replicate, by Arima Genomics using a modified version of the Arima-HiC kit. Briefly, the current Arima-HiC kit (P/N: A510008) utilizes 2 restriction enzymes for simultaneous chromatin digestion. In the modified protocol, 4 restriction enzymes were deployed to enable more uniform per base coverage of the genome while maintaining the highest long-range contiguity signal, thereby benefiting analyses such as base polishing, scaffolding, and phasing. After the modified chromatin digestion, digested ends were labelled, proximally ligated, and then proximally-ligated DNA was purified. After the Arima-HiC protocol, Illumina-compatible sequencing libraries were prepared by first shearing purified Arima-HiC proximally-ligated DNA and then size-selecting DNA fragments using SPRI beads. The size-selected fragments containing ligation junctions were enriched using Enrichment Beads provided in the Arima-HiC kit, and converted into Illumina-compatible sequencing libraries using the Swift Accel-NGS 2S Plus kit (P/N: 21024) reagents. After adapter ligation, DNA was PCR amplified and purified using SPRI beads. The purified DNA underwent standard QC (qPCR and Bioanalyzer) and sequenced on the HiSeq X following manufacturer's protocols.

#### Supplementary Note 3. Assembly and chromosome X finishing

##### Nanopore and PacBio whole-genome assembly

Canu 1.7.1 was used for analysis with the parameters `genomeSize=3.1g corMhapSensitivity=normal ovlMerThreshold=500 correctedErrorRate=0.085 trimReadsCoverage=2 trimReadsOverlap=500 -pacbio-raw` for both data types (Nanopore and PacBio). The X was selected for finishing based on an earlier assembly using the same PacBio data but including only Oxford Nanopore data generated on or before 2018/08/29. Reads were mapped to the assembly using Minimap2 with parameters `-ax map-ont` to identify those spanning gaps and not included in the assembly. The X chromosome, excluding the centromere, was polished using one round of Medaka using only reads assigned by the assembler to the X chromosome. Arrow [2] was run using the ArrowGrid pipeline available at https://github.com/skoren/ArrowGrid using only the P6-C4 chemistry data listed here: https://github.com/nanopore-wgs-consortium/CHM13. The default alignment identity was changed from 0.75 to 0.85. The full assembly, excluding the X centromere, was polished using Nanopolish v0.11.0 using the pipeline available at https://github.com/skoren/NanoGrid. Reads were mapped using Minimap2 with the options `-ax map-ont`. Nanopolish used options `variants --methylation-aware=cpg --consensus -min-candidate-frequency 0.01 --fix-homopolymers`. Arrow v2.2.2 from SMRTlink 6.0.0.47841 was run on the full assembly, again excluding the centromere, with the mapping identity increased to 0.85 `--minAccuracy=0.85 --minLength=5000 --minAnchorSize=12 --maxDivergence=30 --concordant --algorithm=blasr --algorithmOptions=--useQuality --maxHits=1 --hitPolicy=random --seed=1` and additional parameters `-x 10 -q 0 -X120 -v --algorithm=arrow`.

##### 10x Genomics whole-genome assembly and validation

The 10x data was assembled with Supernova v2.1.1 using the command `run --maxreads=all --id=CHM13 --fastqs=Chromium`. This resulted in a 2.95 Gbp assembly with a contig NG50 of 209.7 kbp and scaffold NG50 of 38.5 Mbp for pseudohaplotype 1 and a 2.95 Gbp assembly with a contig NG50 of 209.7 kbp and scaffold NG50 of 38.5 Mbp for pseudohaplotype 2.

Prior to optical mapping, 10x Genomics / Illumina data was mapped to the full assembly using Long Ranger v2.2.2 with the options `longranger align --jobmode=slurm --localcores=32 --localmem=60 --maxjobs=500 --jobinterval=5000 --disable-ui --nopreflight`. Any regions with ≥10-fold coverage were marked as supported. Adjacent supported regions were merged if they were within 500 bp of each other. This list of supported regions was inverted and any unsupported regions within 2 kbp of each other were merged. Finally, the assembly was split at any low-coverage region ≥50 kbp.

##### Bionano optical map assembly and scaffolding

The raw data was assembled with the Bionano Solve data analysis software. This software generated whole genome map assemblies, along with alignments to the reference sequences. In this case, the CHM13 assembly was aligned with CHM13 optical map. After breaking potential mis-assemblies identified by the 10x data, hybrid scaffolding was run using the optical map data using the command `hybridScaffold.pl -n $ASM -b DLE1.cmap -c hybridScaffold_DLE1_config.xml -r avx/RefAligner -B 2 -N 2 -f -o $PWD/scaffold`.

**Hi-C analysis**

Hi-C read mapping heatmap was generated using Juicer v1.5.6 available from https://github.com/VGP/vgp-assembly/tree/master/pipeline/juicer. The restriction site position was indexed with `python juicer-1.5.6/misc/generate_site_positions.py MboI asm asm.fasta` and .hic files were generated with default options ``juicer.sh -z `pwd`/reference/asm.fasta -y `pwd`/reference/asm_MboI.txt -D /usr/local/apps/juicer/juicer-1.5.6/ -d `pwd` -p `pwd`/reference/chr.sizes``. The maps were visualized with Juicebox v1.8.8.

**Chromosome X validation and fixes**

The assembled optical map was used to call high-confidence structural variants on the entire assembly, including the candidate X chromosome. This identified four structural variants (Supplementary Table 4). These SVs were confirmed by discordantly mapping reads later identified in the rel2 ultra-long dataset. To correct these assembly errors, reads over 100 kb with breaks near the variant site were extracted for each SV, making four sets of reads. Each read set was then assembled separately with default parameters by both Canu 1.8 and Flye 2.4 [3,4]. The two assemblers had good agreement and the Flye contigs were aligned to the chromosome X draft and used as patches to replace the incorrect sequence in the original assembly. The patched assembly was once again validated by the optical map, which now reported no discrepancies. PacBio HiFi reads were aligned to the X chromosome and potential repeat collapses identified using a previously described method [5]. This analysis identified the GAGE locus (48.7–48.9 Mbp), cenX (57–61 Mbp), 122 kb segmental duplication containing CXorf49 gene copies (69.5–71.2 Mbp), and CT45 (138.6–139.7 Mbp) as regions of potential collapse (Supplemental Figures 9 and 10). Manual inspection as well as optical map support suggested these regions were not typical repeat collapses, but residual consensus errors due to uneven polishing of large repeat arrays, which were later resolved using a novel polishing strategy as described below.

**Chromosome X long-read polishing**

Unique *k*-mers were identified as those having a copy number in the Illumina read set roughly equal to the expected depth of coverage (between 5 and 58, Supplemental Figure 6) using Meryl[6] from Canu snapshot v1.8 +298 changes (r9508 aab8e5dc15c6b20addccd809c2cc6a62c1fa9c46). In brief, *k*-mers were counted with `meryl count k=21 output 10x.meryl $FASTQ` and filtered with `meryl greater-than 5 output 10x.gt5.meryl 10x.meryl` and `meryl less-than 58 output 10x.gt5.lt58.meryl 10x.gt5.meryl`. Those *k*-mers having both the expected copy number in the 10x data and occuring once in the assembled genome were selected as putative unique markers. `meryl equal-to 1 output asm_1.meryl [ count k=21` asm.fasta ] was run to collect single-copy kmers in the assembly, and it was intersected with meryl intersect output 10x_asm_single.meryl 10x.gt5.lt58.meryl asm_1.meryl. Reads were mapped using Minimap2 v2.71-941 with the parameters -N 50 -r 10000 -ax map-ont. These parameters increase the number of candidate sites reported for a read and tolerate larger gaps within a read without breaking to better allow correction of larger indels in repeat arrays. The Minimap2 alignments were converted to sequence, replacing any mis-matched or missing bases in the read with Ns, and these sequences were scored using the unique markers and placed in the location maximizing the unique marker matches. This generated a new SAM file with all uniquely placed reads assigned a Phred mapping quality (MQ) value of 60. This SAM was filtered to exclude short CIGAR strings (<50 kb for Nanopore, <10 kb for PacBio), and those below a minimum length / identity threshold (25 kb at 75% identity for Nanopore and 5 kb at 75% identity for PacBio). Racon used the parameters -w 5000 -e 0.2. Nanopolish v0.11.0 ran with minimap2 -ax map-ont -N 50 -r 10000 for mapping and nanopolish variants --methylation-aware=cpg --consensus --min-candidate-frequency 0.01 --fix-homopolymers for consensus. Arrow v2.2.2 ran with minimap2 -ax map-pb -N 50 -r 1000 for mapping and -x 10 -q 0 -X120 -v --algorithm=arrow for consensus.

**Whole-genome short-read polishing**

The 10x data was mapped to the scaffolded and polished assembly using Long Ranger v2.2.2 and the options longranger align --jobmode=slurm --localcores=32 --localmem=60 --maxjobs=500 --jobinterval=5000 --disable-ui --nopreflight. FreeBayes was used to call variants with the command freebayes -I -F 0.5 -m 50 --min-alternate-total 5 --min-coverage 10 --max-coverage 100 --read-snp-limit 5 --read-mismatch-limit 5, which enforces a conservative minimum MQ of 50 and only corrects indels supported by more than half of the Illumina reads. This was repeated for two rounds.

**Assembly quality estimation**

We estimated final assembly QV and completeness using previously sequenced CHM13 BACs targeting segmental duplications (VMRC59 library), as well as concordance with the 10x Genomics data. All nucleotide sequences matching VMRC59 with “complete” in the name were downloaded from NCBI. This gave a total of 341 complete BACs. The BACs were mapped with minimap2 with the command minimap2 --secondary=no -ax asm20 -r 2000 and evaluated using the pipeline available from https://github.com/skoren/bacValidation. For 10x Genomics, both Supernova haplotypes were combined and a BAC was considered resolved if either pseudo-haplotype assembly captured it. Out of these 341 BACs, 280 mapped over 99.5% of their length to our CHM13 assembly, which compares favorably to previous assemblies (main text, Table 1). The identity of all BACs mapping over 99.5% of their length was also high for our assembly at 99.98% (Q37.04) median/99.80% (Q27.05) mean vs 99.98% (Q37.32)/99.72% (Q25.60) for PacBio CLR w/ FALCON + Quiver + Pilon, 99.98% (Q36.86)/99.76% (Q26.25) for PacBio HiFi w/ Canu, 99.97% (Q35.97)/99.86% (Q28.45) for 10x Genomics w/ Supernova, and 99.73% (Q25.70)/99.48% (Q22.87) for GRCh38. Using the 31 unique BACs, the identities increase further to 99.99% (Q42.29) median/99.98% (Q36.51) mean vs 99.99% (Q42.68)/99.98% (Q36.75) for PacBio CLR FALCON + Quiver + Pilon, 99.99% (Q44.95)/99.98% (Q37.28) for PacBio HiFi w/ Canu, 99.98% (Q38.12)/99.90% (Q30.30) for 10x Genomics w/ Supernova, and 99.77% (Q26.34)/99.72% (Q25.60) for GRCh38.

Unique BACs were defined as those originating from regions at least 10 kb away from the nearest known segmental duplication. These accessions are: AC275297.1, AC275300.1, AC270133.1, AC270118.1, AC270136.1, AC275290.1, AC279018.1, AC270119.1, AC278482.1, AC275298.1, AC270134.1, AC279070.1, AC270238.1, AC270117.1, AC270132.1, AC270122.1, AC270137.1, AC270115.1, AC275304.1, AC270145.1, AC270121.1, AC278741.1, AC275291.1, AC275285.1, AC270135.1, AC270131.1, AC278929.1, AC275301.1, AC270146.1, AC275305.1, AC270120.1.

We also estimated assembly quality by measuring concordance of the consensus sequence with mapped 10x Genomics / Illumina data. Using the 10x mapping procedure described above, the bam file was filtered for mapping quality >20 with `samtools view -hb -q20`. Variants were called using the command `freebayes --skip-coverage 648 asm.bam -v asm.bayes.vcf -f asm.fasta`, excluding regions with excessive read coverage (12 x mean = 648). Calls genotyped as 0/1 (with support for the assembly allele) were filtered out and the total bases changed (added/deleted/substituted) *B* was summed. Total bases with at least 3-fold and less than 648-fold coverage, *T,* were also tabulated and the QV computed as $-10\ log_{10}(B/T)$, resulting in an average consensus quality estimate of 99.9896% (Q39.83). Note that these FreeBayes parameters are more aggressive and will call more variants than those used for polishing (e.g. the FreeBayes polishing only corrects indels), but this validation is still somewhat circular and we view the BAC validation as more reliable. Using the same criteria, measuring the QV on the X chromosome resulted in 99.9953% (Q43.31).

#### Supplementary Note 4. Structural variant analysis

To compare our CHM13 assembly to GRCh38 as a reference for calling structural variation, contigs from several human assemblies were aligned to each of the two references with MUMmer version 3.23 [7], and structural variants were called using Assemblytics [8]. The four assemblies shown in Supplementary Figure 3 are: (1) the maternal haplotype of NA12878 [6] (2) TrioCanu assemblies of the maternal haplotypes of the Puerto Rican son HG00733 and the Yoruba son NA19240 [9] and (3) a haplotype-phased assembly of a Korean individual [10]. When aligned to GRCh38, the four assemblies yield the following numbers of insertions/deletions: NA12878: 6785/4265, HG00733: 7861/4667, NA19240: 7993/5886, and AK1: 8176/5781. Aligned to the CHM13 assembly, the four assemblies give the following number of insertions/deletions: NA12878: 4127/4345, HG00733: 5018/4897, NA19240: 5709/6578, and AK1: 5656/6113. This excess of insertion calls with respect to GRCh38 exists across a wide size range, and is absent in calls against CHM13.

#### Supplementary Note 5. Determination of copy number of repetitive regions using droplet digital PCR (ddPCR)

Genomic DNA was isolated using DNeasy Blood & Tissue Kit (Qiagen). DNA was quantified using Qubit Fluorometer with Qubit dsDNA HS Assay (Invitrogen). 20uL reaction were performed with 1 ng of gDNA, except for DXZ1 which was run with 0.1 ng of gDNA. Primers and restriction enzymes are listed in the supplemental table. EvaGreen ddPCR reactions were performed using the manufacturer's protocol (Bio-Rad). Mastermixes were simultaneously prepared for HPRT1 and the gene of interest which were then incubated for 15 minutes to allow for restriction digest. Statistics were performed using the confidence interval calculated by the Quantasoft software and applying it to Taylor's expansion.

| **Chromosome region** | **Forward Primer (5'-3')** | **Reverse primer (5'-3')** | **Restriction enzyme** |
|---|---|---|---|
| CT45 | CATCAGCCATGGTGGAGTAT | TGCGGTGTTTCCCTGTT | HaeIII |
| CT47 | GAGATCGGACCCGATGATTC | CCAGTAAATCTCCCACCC | AluI |
| DXZ1 | TGATAGCGCAGCTTTGACAC | TTCCAACACAGTCCTCCA | HaeIII |
| DXZ4 | CACTTCTACCACCACGAGTAA | GGGATGACATTCAACTGGGA | AluI |
| GAGE | GTAACGGAGGTCGTGGATTA | CGCACTGAGAATAAGGGAG | AluI |
| **Reference Gene** | **Forward Primer (5'-3')** | **Reverse primer (5'-3')** | **Restriction enzyme** |
| HPRT1 | AAGGTGCTGGTCTCCTTTAC | GCACCAATGATTCTCTCCCT | AluI |

#### Supplementary Note 6. Chromosome X centromere (DXZ1) array PFGE Southern analysis

##### Pulsed field gel electrophoresis

Alpha satellite array sizes were estimated by PFGE and Southern blotting using established methods [11,12]. High molecular weight DNA from $10^7$–$10^8$ was embedded in 1% low melting point agarose plugs and digested with restriction enzymes that cut infrequently within alpha satellite DNA, releasing the DXZ1 array as one of a few large fragments. HMW DNA in one-half of an agarose plug was digested overnight with 20U of enzyme and run on 1% agarose gel. *Saccharomyces cerevisiae* and *Hansenula wingei* chromosomes embedded in agarose were used as size standards (Bio-Rad CHEF DNA Size Markers). Gels were run at 3 volts/cm for 50 hours at 14 °C in 1X TAE buffer, using switch times of 250 seconds (initial) – 900 seconds (final). Cell lines containing previously sized DXZ1 arrasy were used as controls [11,12].

**Southern blotting**

After electrophoresis, gels were stained with ethidium bromide and imaged using a UV light source. Gels were rinsed briefly with distilled water, depurinated with 0.25 M HCl for 12 minutes at room temperature, then incubated twice for 15 minutes in denaturing buffer (1.5 M NaCl, 0.5 M NaOH). DNA was transferred to HyBond-N+ membrane (GE Healthcare/Amersham) for 48 hours in fresh denaturing buffer. Dried membranes were UV crosslinked (auto-crosslink setting on Stratagene Stratalinker) before proceeding to hybridization.

A 500 bp fragment (2 micrograms) spanning monomers 9–12 of DXZ1 was generated by PCR [13] and labeled overnight at 37°C with digoxygenin-11-dUTP using DIG High Prime (Sigma-Aldrich). Alternatively, a plasmid containing an entire DXZ1 HOR (2 kbp) was labeled by nick translation with digoxygenin-11-dUTP for 90 minutes at 14 °C. Labeling reactions were purified using either the High Pure PCR purification kit (Roche) or G-50 sephadex columns.

Membranes were pre-hybridized for 30–45 minutes in glass hybridization bottles containing 20 mL ExpressHyb buffer (Clontech) at 63 °C. Pre-hybridization buffer was replaced with 20 mL of fresh ExpressHyb containing 300–400 ng of labeled probe that had been denatured at 95 °C for 10 minutes. The probe was allowed to hybridize to the membrane at 63 °C overnight in a hybridization oven. Membranes were washed at 68 °C twice for 20 minutes in 2X SSC/0.1% sodium dodecyl sulfate (SDS), followed by a single high-stringency wash in 0.2X SSC/0.1% SDS for 15 minutes at 68 °C. Membranes were blocked in 1x Western blocking reagent (Roche) in maleic acid buffer (0.1 M maleic acid, 0.15 M NaCl, pH 7.5) for 45–60 minutes at room temperature, then incubated for 30 minutes in blocking buffer with anti-digoxygenin-alkaline phosphatase (Roche, 1:2000). Chemiluminescent detection was performed using 4–5 mL of CDP-Star ready-to-use reagent (Tropix). Membranes were imaged on a G:Box using GeneSys software (Syngene) for direct image analysis. Images were adjusted (leveled to curves) and labeled in Adobe Photoshop.

#### Supplementary Note 7. Chromosome X centromere (DXZ1) CRISPR-Cas9 duplex sequencing

**DXZ1 CRISPR-Cas9 in vitro digestion**

CRISPR-DS was performed as previously described [14] for a single sample (CHM13). Briefly, we designed the following guide RNA sequences to excise the DXZ1 centromeric satellite DNA: GAGGGCTTTGAGGCCTGTGGTGG and GTTCCTTCCTATACGACCGTAGG. 30nM of gRNAs were incubated with Cas9 nuclease at 25 °C for 10 min. We used a 0.5X ratio of AMPure beads to size select for the excised DNA fragments. Then the fragments were A-tailed and ligated to adapters including a 10 bp random double-stranded molecular tag (TwinStrand Biosciences) using the NEB kit as described [15]. The ligated DNA was amplified using KAPA Real-Time Amplification kit with fluorescent standards (KAPA Biosystems). Two xGen Lockdown Probes (IDT) specific to DXZ1 (4 nmole Ultramer DNA Oligo, shown below) were used to perform hybridization capture as previously reported with minor modifications [14]. The lockdown probes were pooled in equimolar amounts and diluted to 0.75 pmol/µL in low TE (0.1 mM EDTA).

/5Biosg/GAAACGACTTTGTGAGGATGGCATTCAACTCATGGAGTTGAACAATCCTATTGATA
GAGCAGATTGGAATCACTCTTTTTGTAGAATCTGCAAATGGAGATTTGGACTGCTTTGAGG
CCT

/5Biosg/GAGGCCTGTGGTGGAAAAGGAAATATCTTCACATAAAAACTAGATAGAAACACTCT
GAGAAAGTTCTTCATGATGAATGCATTTAACTCGCAGAGATGAACCTGCCTTTGAGAGTTCA
GG

The CHM13 sample was quantified using the Qubit dsDNA HS Assay Kit, diluted, and pooled for sequencing. The library was sequenced on the MiSeq Illumina platform using a v3 600 cycle kit (Illumina), as specified by the manufacturer. Analysis was performed as previously described [15] using software available: https://github.com/risqueslab

#### Supplemental Figures

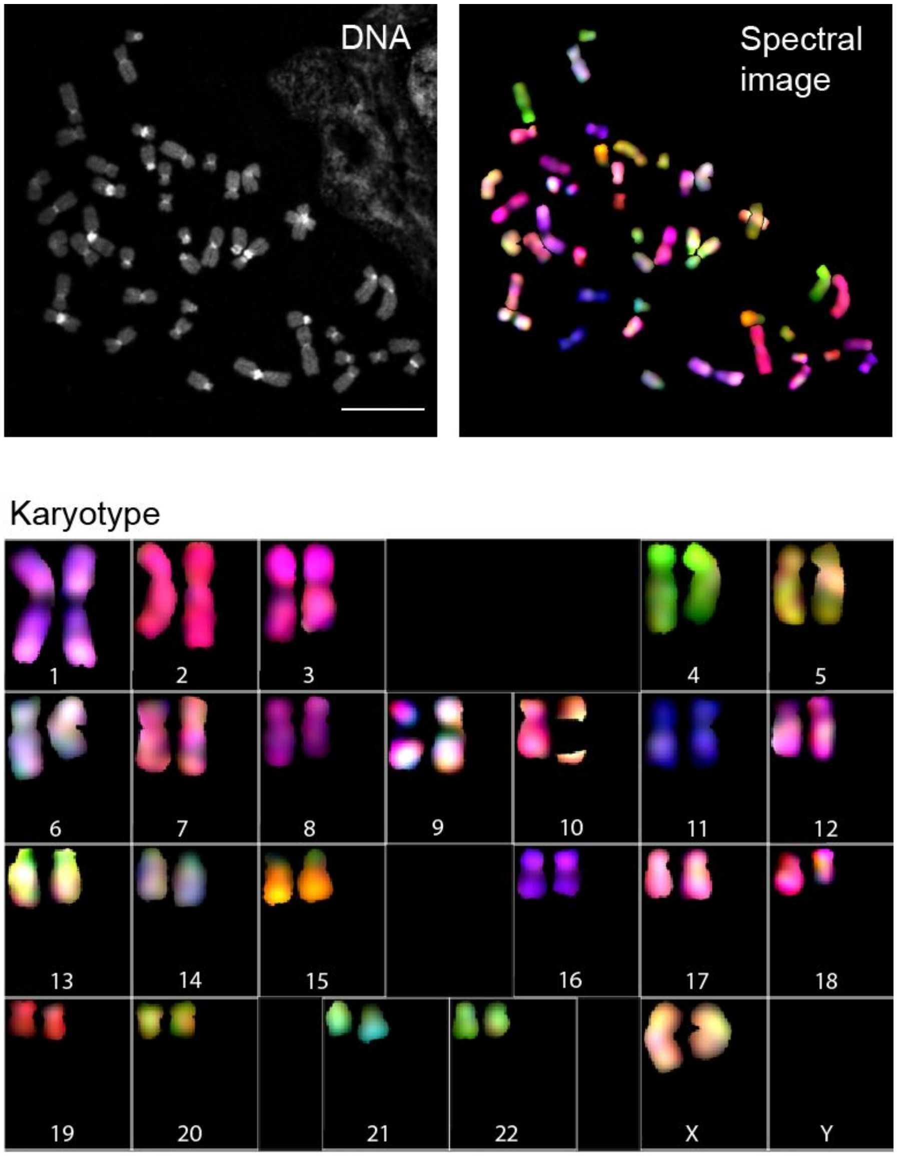

**Supplemental Figure 1.** Chromosomes and karyotype of CHM13 cell line at passage 10. Mitotic metaphase spreads were prepared from cells treated with colcemid and processed as detailed in Materials and Methods. Spectral karyotyping analysis demonstrated normal 46,XX karyotype. Representative karyotype is shown from one of ten spreads analyzed. Bar, 10 μm.

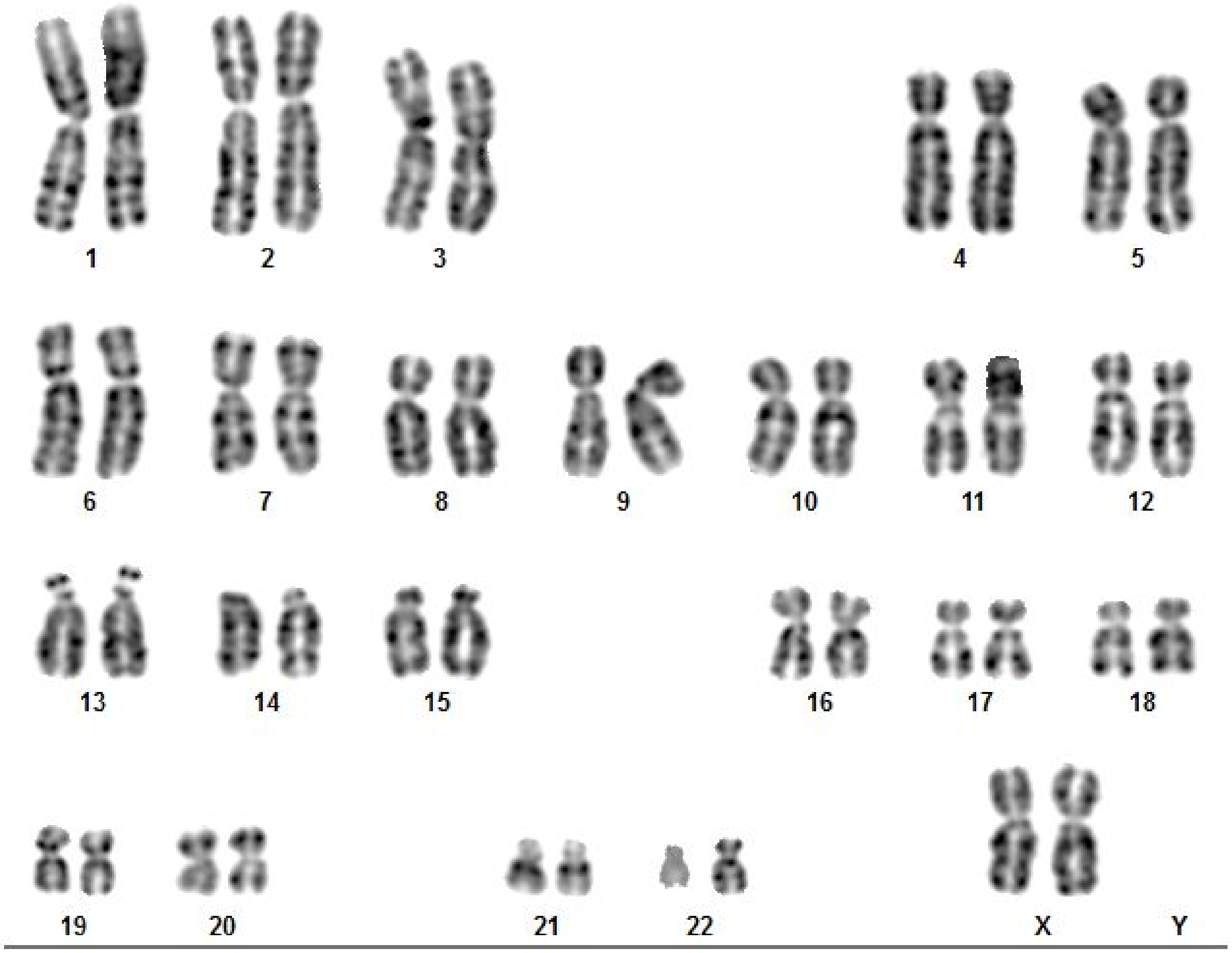

**Supplemental Figure 2.** CHM13 G-banding karyotype. A total of 20 CHM13 metaphase spreads were characterized and all showed a normal 46, XX female karyotype, as shown.

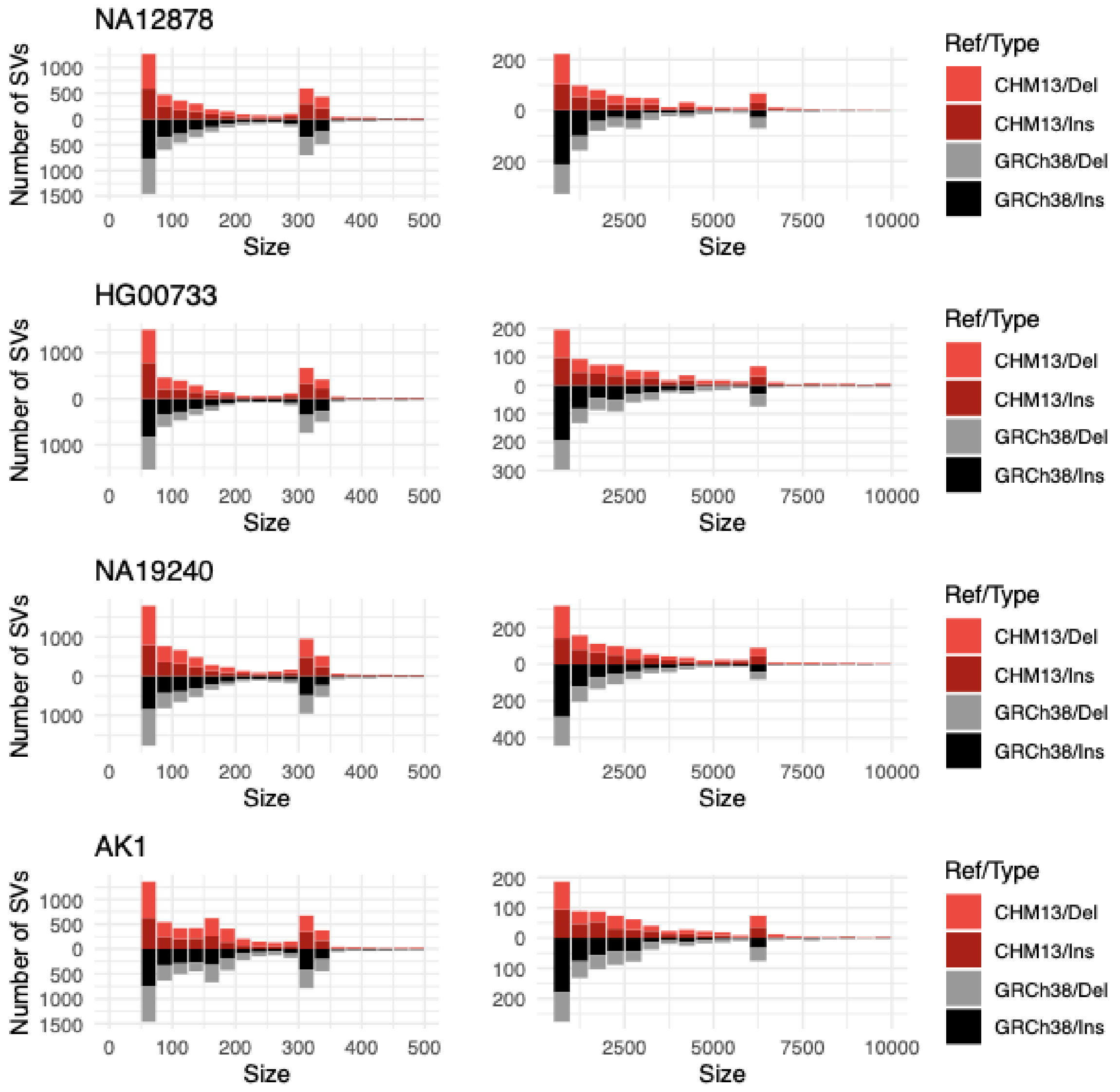

**Supplemental Figure 3.** Results of using CHM13 as a reference when describing structural variation. Assemblytics large insertion and deletion calls for four long read assemblies with respect to CHM13 (in dark red/red) and GRCh38 (in black/gray). Using CHM13 as a reference yields balanced counts of insertions and deletions, while an excess of insertion calls is observed when using GRCh38, suggesting a probable deletion bias in GRCh38.

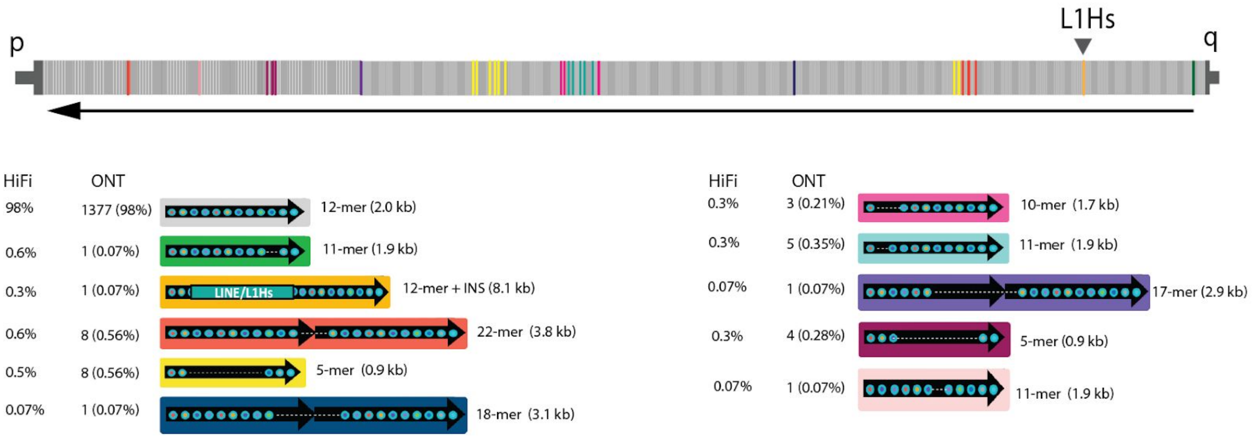

**Supplemental Figure 4.** Comparisons with DXZ1 higher-order repeat variant frequency in the nanopore UL data with high-fidelity (HiFi) long-read pacbio data [16] were highly concordant. DXZ1 repeat unit variants were predicted in the HiFi dataset using alpha-CENTAURI [17] DXZ1 repeat unit, shown as arrows, are composed of 12 smaller ~171 bp repeats (indicated as small circles within the arrow). Changes from the canonical repeat unit are indicated with a dashed line and each structural variant marks a color, and its positioning within the array assembly is indicated (ordered p-arm to q-arm) above.

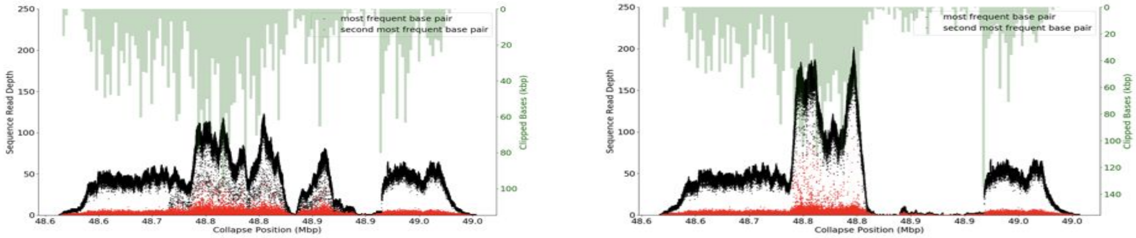

**Supplemental Figure 5.** The initial Canu assembly of the GAGE locus (left) was further corrupted as a result of running standard long-read polishing pipelines (Arrow and Nanopolish, right). Black dots indicate coverage of the primary allele and red dots indicate coverage of the secondary allele based on mapping of PacBio CLR data. The CHM13 genome is effectively haploid so only one allele is expected, and regions of low coverage or increased secondary allele frequency indicate low-quality regions or potential repeat collapses. Due to mis-mapping of reads during the polishing process, allele coverage becomes even less uniform after polishing. A modified polishing process, using the unique *k*-mer strategy, corrects this effect (SFig 9,10).

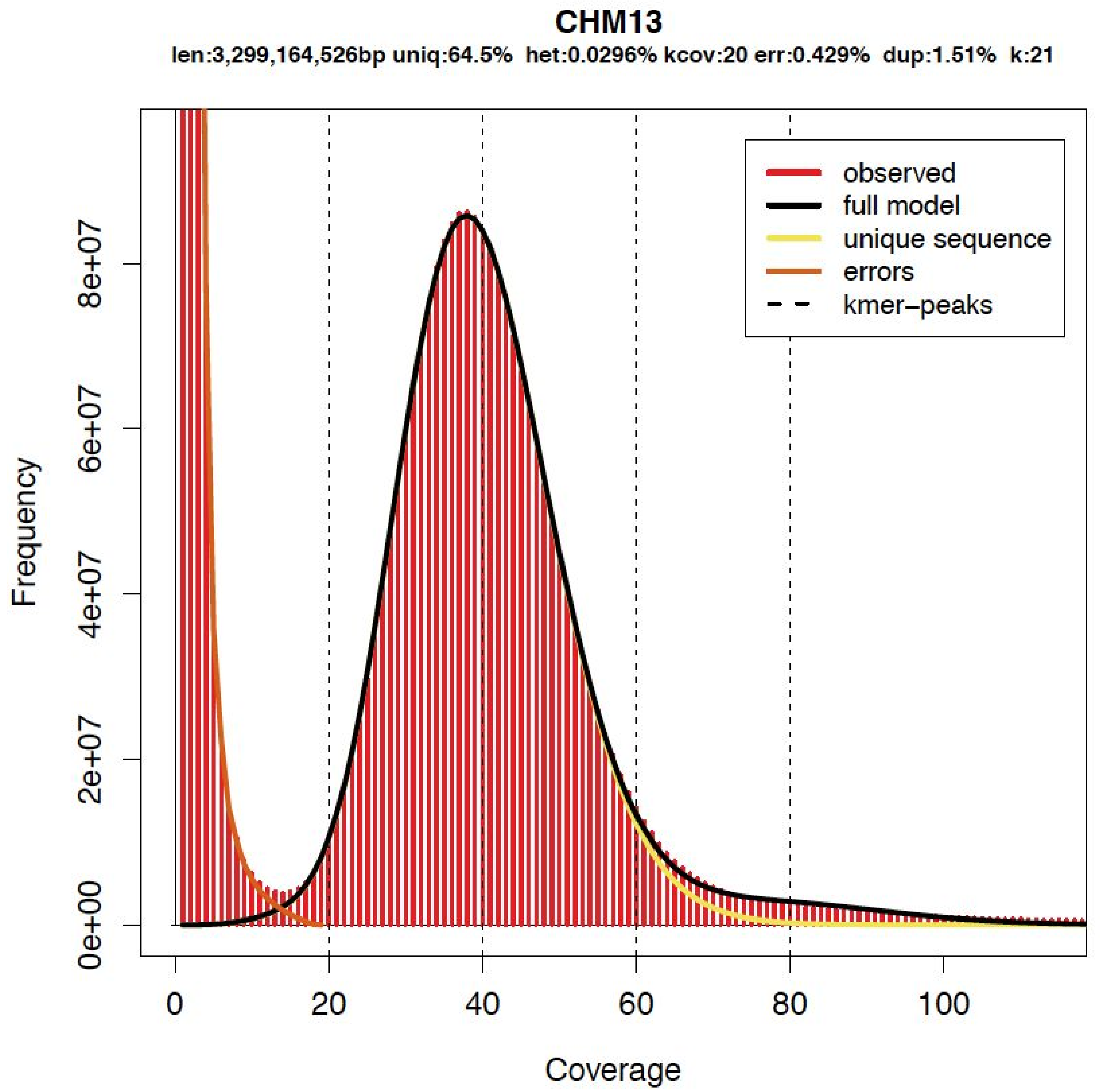

**Supplemental Figure 6.** 21-mer distribution from the 10x Genomics reads. 21-mers were collected with Meryl and the plot was generated with GenomeScope1.0 [18] to visualize and confirm the haploid nature of CHM13 and genome size (len). k-mers with counts between 5 and 58 (inclusive) were used as unique markers when polishing the X chromosome.

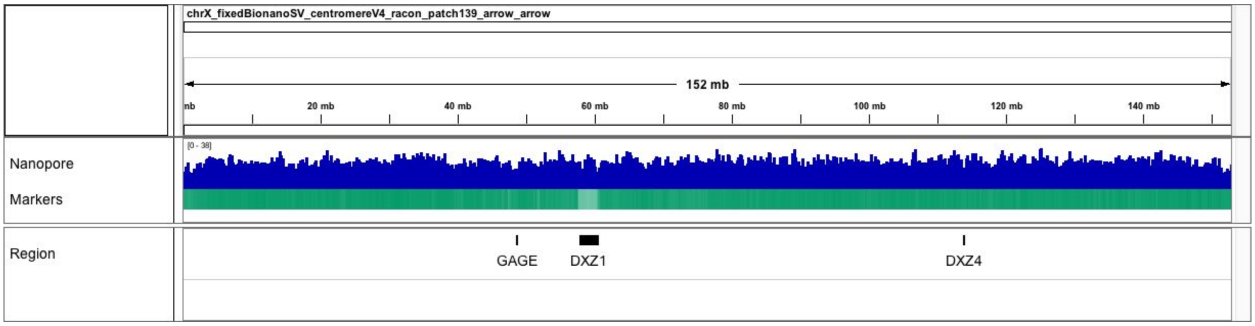

**Supplemental Figure 7.** Mapped nanopore reads show uniform coverage across the complete X chromosome. Reads were filtered using the same unique marker based filtering as for polishing. Marker density is shown below the read alignments.

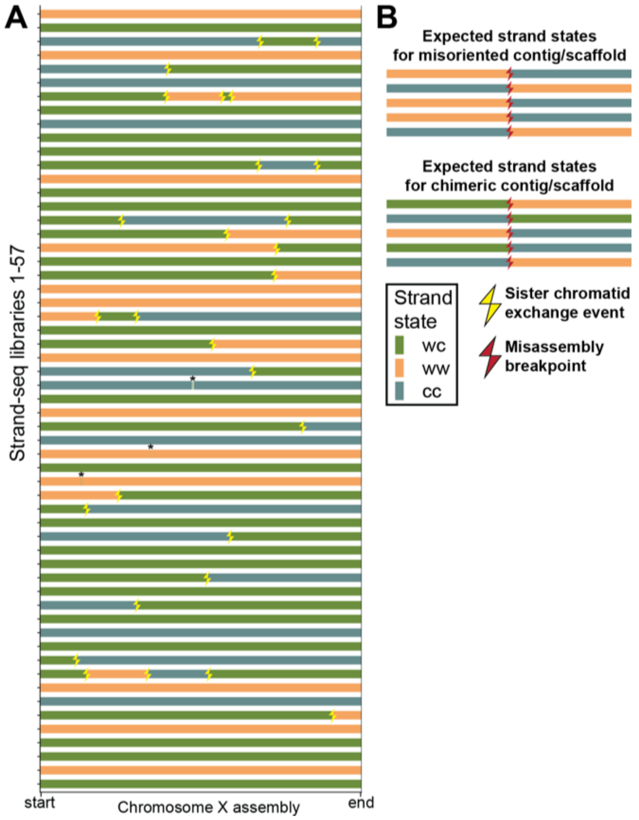

**Supplemental Figure 8.** Strand-seq validation of the chromosome X assembly. Strand-seq sequences only single template strands from each homologous chromosome. Sequencing reads originating from such single stranded DNA posses directionality, a feature that can be used to assess a long range contiguity of individual homologs[19,20]. Based on the inheritance of single stranded DNA we distinguish 3 possible strand states: WW – both homologs inherited Watson template strand, CC – both homologs inherited Watson template strand and WC – one homolog inherited Watson and the other Crick template strand. By tracking changes in strand states along each chromosome we are able to pinpoint locations of recurrent strand state changes that are indicative of a genome misassembly. (A) We have analyzed in total 57 Strand-seq libraries and mapped 28 localized strand state changes. These strand state changes are randomly distributed along chromosome X assembly and therefore are indicative of a double-strand-break that occurred during DNA replication instead of real genome misassembly. Such breaks are usually repaired by available sister chromatid and therefore often result in change in strand directionality. Black asterisks show small localized strand state changes. Such events are either caused by a noisy reads inherent to Strand-seq library preparation or two double-strand-breaks that occurred very close to each other. (B) Because it is unlikely for a double-strand-break to occur at exactly the same position in multiple single cells, a real genome misassembly is visible in Strand-seq data as a recurrent change in strand state at the same position in a given contig or scaffold. None of these signatures was observed in the CHM13 chromosome X assembly.

a

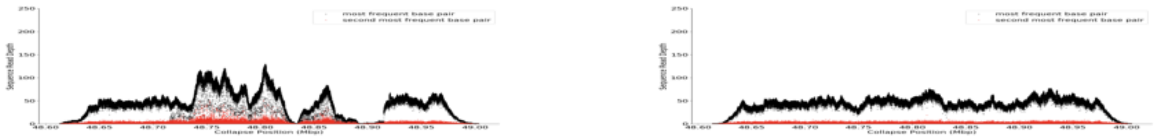

b

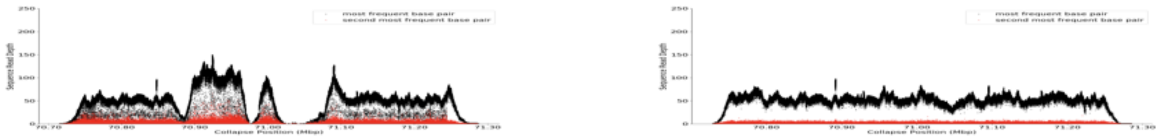

c

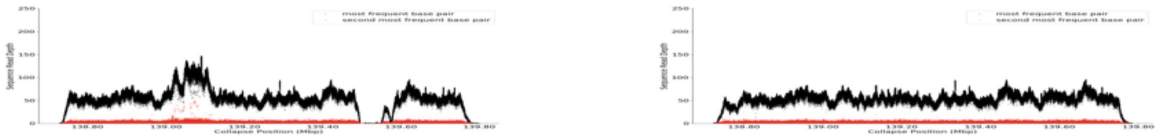

d

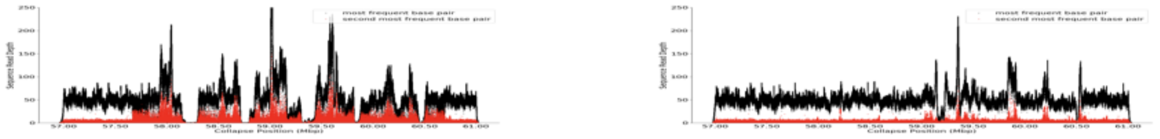

**Supplemental Figure 9.** Final evaluation of repeat base-level quality was determined by mapping of PacBio CLR data. Dots indicate coverage depth of the primary (black) and secondary (red) alleles recovered from mapped PacBio CLR sequences. The left-side plots indicate assembly evaluated before the polishing process. The right-side plots show the same regions after using unique marker-assisted polishing (racon, 2 rounds of nanpolish, 2 rounds of arrow, 2 rounds of 10x (SNote 3). The regions shown are a) GAGE locus (48.6-49 Mbp), b) 70.8-71.3 Mbp, c), 138.6-139.7 Mbp, and d) cenX (57-61 Mbp).

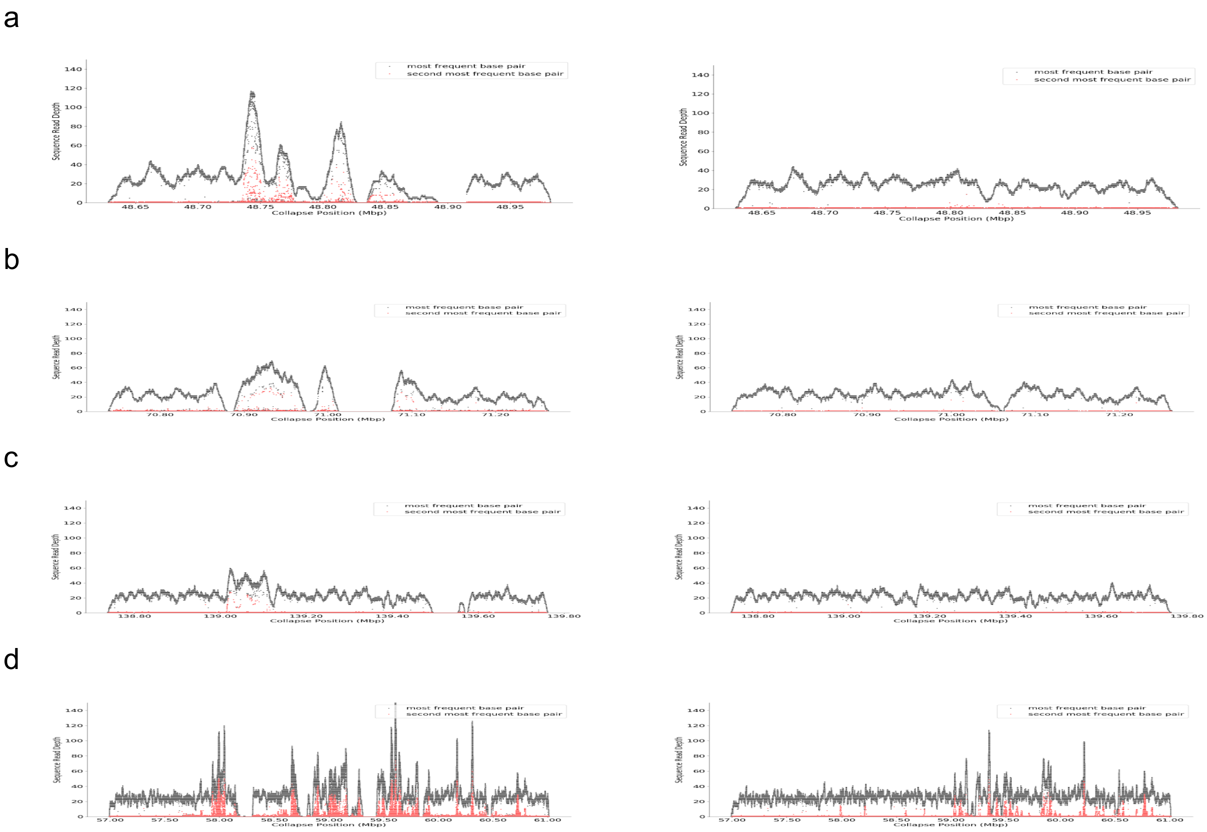

**Supplemental Figure 10.** Evaluation as in SFig 9 using HiFi rather than CLR PacBio data.

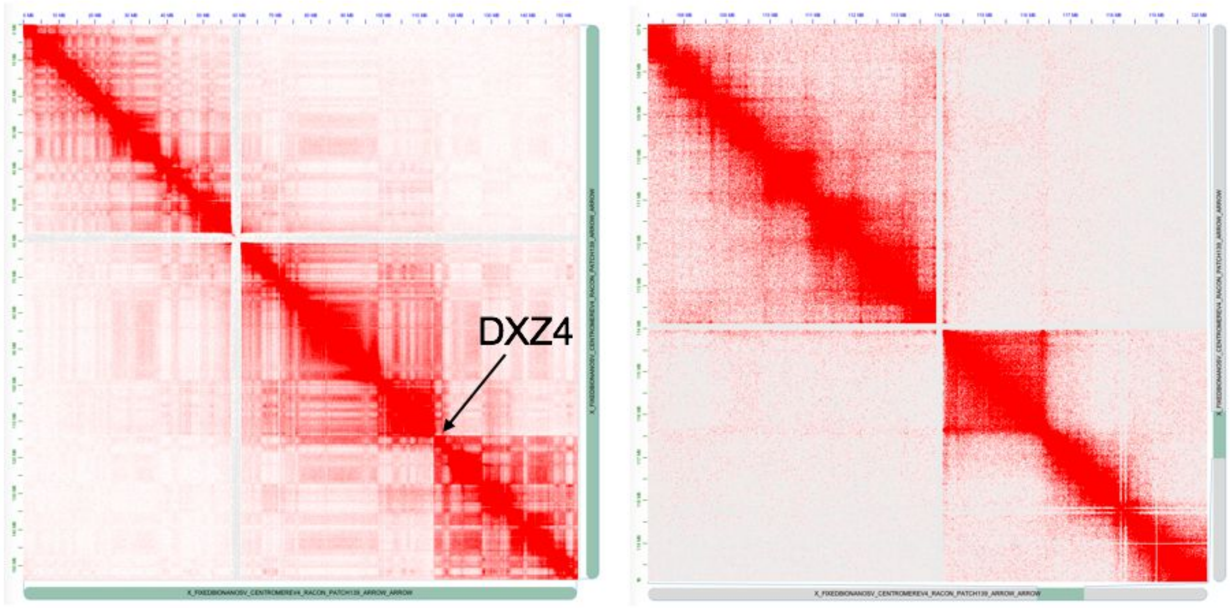

**Supplemental Figure 11.** Hi-C read mapping to the chromosome X assembly. The whole X is shown on the left, and the right is zoomed on the DXZ4 locus. The heatmap shows clear boundaries around DXZ4, indicating 2 large superdomains separated by DXZ4.

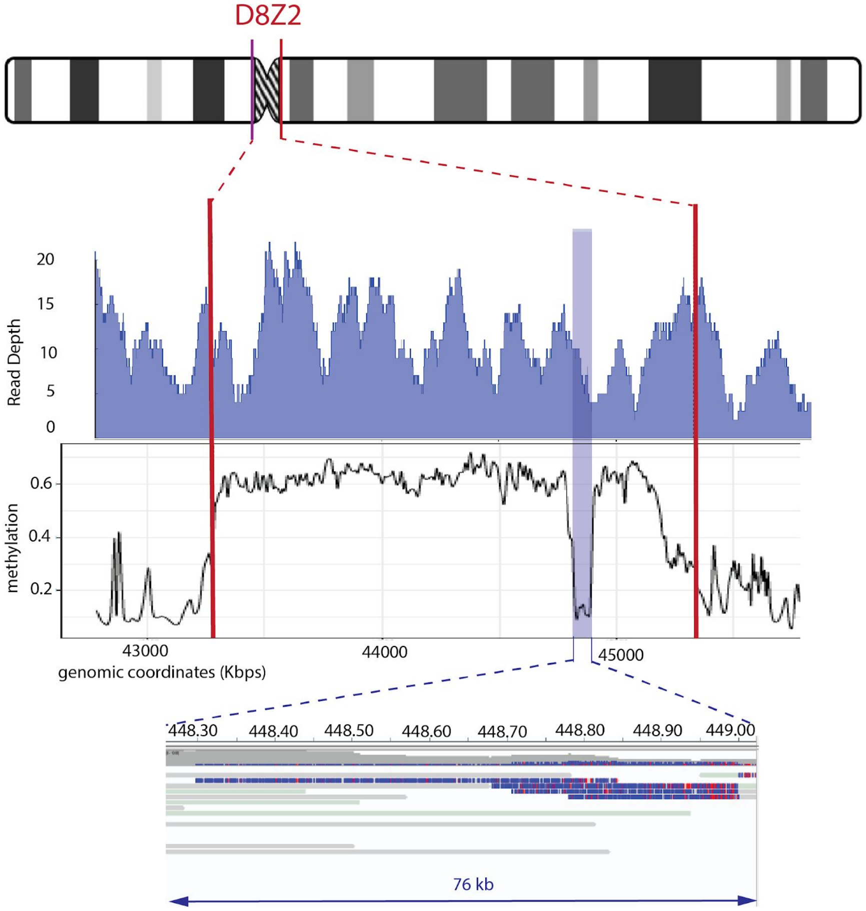

**Supplemental Figure 12.** Methylation estimates across centromeric satellite array assembly on chromosome 8 (D8Z2) (chr8:43,281,085-45,333,062). Methylated values were calculated by smoothing frequency data with a window size of 500 nucleotides. Read coverage shown relies on our unique-anchor mapping and the presence of at least one high-quality methylation call on the read |log-likelihood| > 2.5. Similar to our previous methylation analysis on chromosome X centromeric satellite array (DXZ1), we observe an unmethylated region (~75 kb) in the centromere of chromosome 8 (as shown: chr8:44,830,000-44,900,000).

### Supplemental Tables

| Name | T2T X | T2T WG |
|---|---|---|
| GenesFound | 841 | 19618 |
| GenesFoundPercent | 99.64 | 99.68 |
| TranscriptsFound | 2994 | 83332 |
| TranscriptsFoundPercent | 99.87 | 99.82 |

| | | |
|---|---|---|
| FullmRNACoverage | 2628 | 71684 |
| FullmRNACoveragePercent | 87.66 | 85.87 |
| FullCDSCoverage | 2788 | 77114 |
| FullCDSCoveragePercent | 93.00 | 92.37 |
| TranscriptsWithFrameshift | 19 | 334 |
| TranscriptsWithFrameshiftPercent | 0.63 | 0.40 |
| TranscriptsWithOriginalIntrons | 2771 | 77927 |
| TranscriptsWithOriginalIntronsPercent | 92.43 | 93.35 |
| TranscriptsWithFullCDSCoverage | 2788 | 77114 |
| TranscriptsWithFullCDSCoveragePercent | 93.00 | 92.37 |
| TranscriptsWithFullCDSCoverageAndNoFrameshifts | 2788 | 77101 |
| TranscriptsWithFullCDSCoverageAndNoFrameshiftsPercent | 93.00 | 92.36 |
| TranscriptsWithFullCDSCoverageAndNoFrameshiftsAndOriginalIntrons | 2711 | 76632 |
| TranscriptsWithFullCDSCoverageAndNoFrameshiftsAndOriginalIntronsPercent | 90.43 | 91.80 |
| GenesWithFrameshift | 9 | 170 |
| GenesWithFrameshiftPercent | 1.07 | 0.86 |
| GenesWithOriginalIntrons | 803 | 18490 |
| GenesWithOriginalIntronsPercent | 95.14 | 93.95 |
| GenesWithFullCDSCoverage | 794 | 18314 |
| GenesWithFullCDSCoveragePercent | 94.08 | 93.06 |
| GenesWithFullCDSCoverageAndNoFrameshifts | 796 | 18355 |
| GenesWithFullCDSCoverageAndNoFrameshiftsPercent | 94.31 | 93.27 |
| GenesWithFullCDSCoverageAndNoFrameshiftsAndOriginalIntrons | 788 | 18330 |
| GenesWithFullCDSCoverageAndNoFrameshiftsAndOriginalIntronsPercent | 93.36 | 93.14 |
| MissingGenes | 3 | 62 |
| MissingGenesPercent | 0.36 | 0.32 |

**Supplemental Table 1.** Genome annotation results from the Comparative Annotation Toolkit (CAT) for the CHM13 assembly presented here. Results are provided for both chromosome X and the whole genome.

| Assembly Name | Sample | Assembler | Cov | Instrument / Chemistry | # Ctg | Size (Gbp) | NG50 (Mbp) |
|---|---|---|---|---|---|---|---|
| GCA_000983475.1 | CHM13 | Celera Assembler | 70x | RSII/P5+P6 | 10,430 | 3.00 | 5.35 |
| GCA_000983455.2 | CHM13 | Falcon | 70x | RSII/P5+P6 | 4,961 | 2.94 | 9.85 |
| GCA_001015385.3 | CHM13 | Celera Assembler | 70x | RSII/P5+P6 | 12,091 | 3.07 | 11.95 |
| GCA_000983465.1 | CHM13 | Celera Assembler | 70x | RSII/P5+P6 | 15,538 | 3.06 | 12.48 |

| | | | | | | | |
|---|---|---|---|---|---|---|---|
| GCA_001015355.1 | CHM13 | Celera Assembler | 70x | RSII/P5+P6 | 11,138 | 3.03 | 19.03 |
| GCA_001307015.1 | CHM1 | Celera Assembler | 120x | RSII/P5+P6 | 5,307 | 3.01 | 25.37 |
| GCA_001297185.2 | CHM1 | Falcon | 60x | RSII P6 | 3,709 | 3.00 | 26.13 |
| GCA_001524155.4 | NA19240 | Falcon + BioNano | 73x | RSII P6 | 2,439 | 2.87 | 28.15 |
| GCA_002884485.1 | CHM13 | Falcon | 76x | RSII P6 | 1,916 | 2.88 | 28.20 |
| GCA_002180035.3 | HG00514 | Falcon + BioNano | 80x | RSII P6 | 2,799 | 2.86 | 29.00 |
| GCA_001420755.1 | CHM1 | Celera Assembler | 120x | RSII/P5+P6 | 2,416 | 2.95 | 29.05 |
| GCA_001420765.1 | CHM1 | Celera Assembler | 120x | RSII/P5+P6 | 3,188 | 2.99 | 32.45 |
| GCA_000001405.28 | GRCh38p13 | N/A | N/A | N/A | 1,590 | 3.11 | 56.41 |
| T2T v0.6 | CHM13 | Canu | 39x + 70x | Oxford GridION/9.4.1 | 590 | 2.93 | 71.7 |

**Supplemental Table 2.** All human genome assemblies in NCBI with contig NG50 >25 Mbp or originating from CHM13. Sequences were downloaded from the FTP site and scaffolds split at 3 consecutive Ns to get contigs. A genome size of 3.0988 was used for computing NG50 for all assemblies. Aside from the Nanopore assembly presented here, all other assemblies in the table were generated using PacBio CLR data. The CHM13 PacBio CLR assembly we compare against in the main text is GCA_002884485.1 which had the highest score for BAC resolution of all CHM13 assemblies tested and incorporated the highest coverage PacBio data.

| Cell line | PFGE DXZ1 Estimation | ddPCR DXZ1 Estimation |
|---|---|---|
| HAP1 | 3.7 Mb | 3.7 Mb |
| t60-12 | 3.0-3.1 Mb | 3.2 Mb |
| HDF | 3.8 Mb | 2.9 Mb |
| LT690 | 1.5 Mb | 1.4 Mb |
| CHM13 | 2.8 Mb | 2.8 Mb |

**Supplemental Table 3.** DXZ1 array estimations for five different cell lines using PFGE and ddPCR. *HPRT1* was used as ddPCR single copy reference gene. PFGE were the result of at least three different runs with several standards.

| Sequence Name | RefStartPos | RefEndPos | Type | Size |
|---|---|---|---|---|
| chrX_bothkpatchedin | 48,733,807 | 48,790,958 | insertion | 124,036 |
| chrX_bothkpatchedin | 70,270,806 | 70,340,885 | deletion | 25,207 |
| chrX_bothkpatchedin | 106,136,920 | 106,142,580 | insertion | 2,961 |

| | | | | |
|---|---|---|---|---|
| chrX_bothkpatchedin | 133,151,139 | 133,220,191 | insertion | 17,489 |

**Supplemental Table 4.** Structural variants identified by BioNano optical map in chromosome X draft. A table displaying coordinates and sizes of SVs identified in the candidate chromosome X draft.
